# Reported Health Benefits of a Vegan Dog Food – A Likert Scale-type Survey of 100 Guardians

**DOI:** 10.1101/2022.05.30.493980

**Authors:** Mike Davies

## Abstract

**Introduction:** There has been a surge in feeding plant-based foods to pets and advocates extol the health benefits of this practice. However, there is a lack of scientific evidence to support health claims for vegan diets in dogs.

**Aims:** This study aimed to quantify perceived health changes by dog guardians following the feeding of a single brand of UK-produced vegan food for a period of 3 to 12 months.

**Methods:** Dog guardians registered as feeding the vegan food for 3 - 12 months were invited to participate in an online Likert Scale-type survey of observations reflecting health status.

**Results:** 100 guardians completed the survey. The vegan food was acceptable (palatable), and appetite and body weight were not adversely affected. Changes, including improvements, were reported in the following areas: body condition score (BCS), activity, faecal consistency, faecal colour, frequency of defaecation, flatus frequency, flatus antisocial smell, coat glossiness, scales in haircoat (dandruff), redness of the skin (erythema, inflammation), crusting of the external ear canals (otitis externa), itchiness (scratching; pruritus), anxiety, aggressive behaviour and coprophagia.

**Conclusions:** This is the first study to quantitatively document guardian reports of specific positive health benefits associated with feeding a UK vegan dog food. Further prospective, randomised, controlled clinical trials are needed to validate and determine the significance of these observations.

## Introduction

In recent years there has been an increase in feeding vegetarian and vegan foods to pets, with the vegan pet food market being valued at $9.6 billion in 2020 and is estimated to reach $16.3 billion by 2030 (Allied Market Research 2022). Proponents of plant-based foods often claim a variety of health benefits without providing scientific evidence to support their claims. A UK-based company (Omni) launched a range of plant-based (vegan) dog foods in April 2021 and noticed that some customers were posting messages online through a feedback service Trustpilot claiming improvements in their pets’ health, particularly skin and gastrointestinal problems. Trustpilot.com is a Danish consumer review website founded in 2007 which hosts reviews of businesses worldwide.

In a recent paper (Knight and others 2022) 336 guardians feeding vegan diets participated in a global online survey and the authors concluded that, “when compared to conventional meat-based or raw meat rations the healthiest and least hazardous dietary choices for dogs, are nutritionally sound vegan diets”. In this study some health issues were mentioned but there were no data on specific changes in health status following a switch to feeding vegan.

Previous published papers have shown that vegan diets labelled as complete and recipes for homemade vegan rations did not comply with AAFCO or FEDIAF Nutritional Recommendations and therefore, contrary to providing health benefits would represent a potential health risk (Dodd and others 2021; Pedrinelli and others 2021; Starzonek and others 2021; Zafalon and others 2019).

One study (Cavanaugh and others 2021) looked at the short-term effects of a vegan diet on amino acid, clinicopathologic, and echocardiographic findings and found no adverse effects and did not detect essential amino acid or taurine deficiency.

All batches of the vegan diet involved in this study are analysed by the University of Nottingham to ensure that they still exceed FEDIAF Guidelines after manufacture. Dog carers who feed them may not be their legal owners and so in this paper they are referred to as guardians.

## Objectives

The aim of this study was to document and quantify any changes in signs associated with health status observed by guardians following a switch to feeding dogs a single brand of complete vegan food for over 3 months.

The aim was to see if there was any improvement or deterioration in observations that may reflect health status:

Acceptability of food when first fed Appetite

Body weight

Body condition score (BCS)

Activity level

Gastrointestinal changes:

a. Faecal consistency
b. Frequency of defaecation
c. Colour of faeces
d. Flatus frequency
e. Flatus antisocial smell

Skin signs

a. Glossiness (shine)
b. Scaly skin (dandruff)
c. Redness (erythema) of the skin (inflammation)
d. Crusting of the external ear canals (otitis externa)
e. Itchiness (scratching; pruritus)

Behaviour

a. Anxiety
b. Aggression
c. Any other behavioural changes

## Hypothesis

The study was designed to acquire data to disprove the hypothesis that “Feeding a vegan diet to dogs does not provide health benefits”

## Ethics

The study was designed to comply with established ethical standards and it was decided that the protocol did not need to be submitted to a formal external review panel. It complied with guidance set out in the Royal College of Veterinary Surgeon’s Ethical Review for Practice-based Research (RCVS 2013) and guidance on ethics in questionnaires published online (Travis 2017)

## Methods

### Data Collection

All data was collected, stored and used in accordance with The Data Protection Act 2018 and the General Data Protection Regulation (GDPR).

### The Dog Food

The plant-based vegan pet food fed to the dogs in this study was formulated by veterinarians and manufactured in the UK. The main ingredients incorporated into the recipes (present at >4% inclusion rate) include: Potato Protein, Pea starch, Soya, Brown Rice, Sweet potato, Dried Brewers Yeast, Oats, Peas, Pea Protein, Carrot and Rapeseed Oil.

### Recruitment of guardians

On 18^th^ March 2022 the company Omni sent individual emails using email marketing software Klaviyo (https://www.klaviyo.com) to 307 randomly selected dog guardians who had been registered as feeding their plant-based food for at least 3 months. They were invited to participate in an online survey to generate basic information and data about perceived changes in health and informed consent was obtained to publish the results afterwards.

The survey went live on 18th March 2022 and was terminated on 25th April 2022 by which time 101 people had completed it.

A second email was sent to 100 respondents later with a few supplementary questions regarding the dog breed, sex and feeding practices before switching to the plant-based food

### The Likert Scale-style Survey

An online Likert Scale-style questionnaire was created and analysed in accordance with established guidelines (Boynton 2004; Sullivan and Artino 2013)

The questions were assessed for clarity, objectivity, appropriateness, avoidance of leading questions and ease of use by veterinarians working with the company. The online questionnaire was created using Typeform (https://www.typeform.com) to collect the data. Apart from the first question on acceptability of the food and the question about colour of the faeces, the questions in the survey were based on a 5-point Likert scale in which the middle option was “No change” and either side were two options to report a slight or great improvement or deterioration in observations that may reflect health status as described and listed in the study objectives:

### Acceptability

Guardians were asked to select one of the following that best described what happened when they first introduced the plant-based food to their dog.

Options:

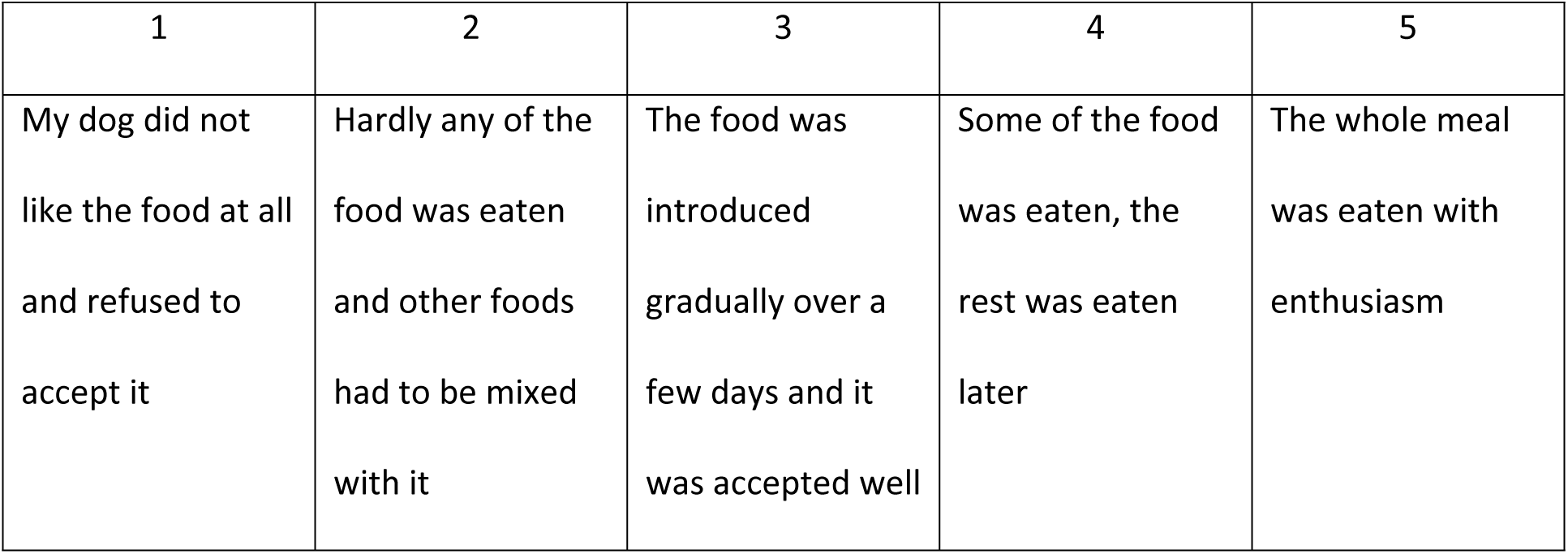

### Appetite

Guardians were asked: How good was your dog’s appetite before you switched to the plant-based food?

After changing to the plant-based food what effect, if any, was there on appetite:

Options:

| 1 | 2 | 3 | 4 | 5 |
| --- | --- | --- | --- | --- |
| Appetite was greatly reduced | Appetite decreased slightly – less hungry | No change in appetite | Appetite increased slightly | Appetite increased a lot with begging behaviour |

### Body Weight

Guardians were asked: After switching to the plant-based food what approximate changes did you notice in body weight?

Options:

| 1 | 2 | 3 | 4 | 5 |
| --- | --- | --- | --- | --- |
| Body weight decreased by more than 10% | Body weight decreased by less than 10% | There was no change in body weight | Body weight increased by less than 10% | Body weight increased by more than 10% |

### Body Condition Score (BCS)

Guardians were asked: If you measure your dog’s Body Condition Score (BCS) using a recognised chart please answer this question (if not please skip this question).

After switching to the plant-based food what change have you noticed in body condition score (BCS)?

Options:

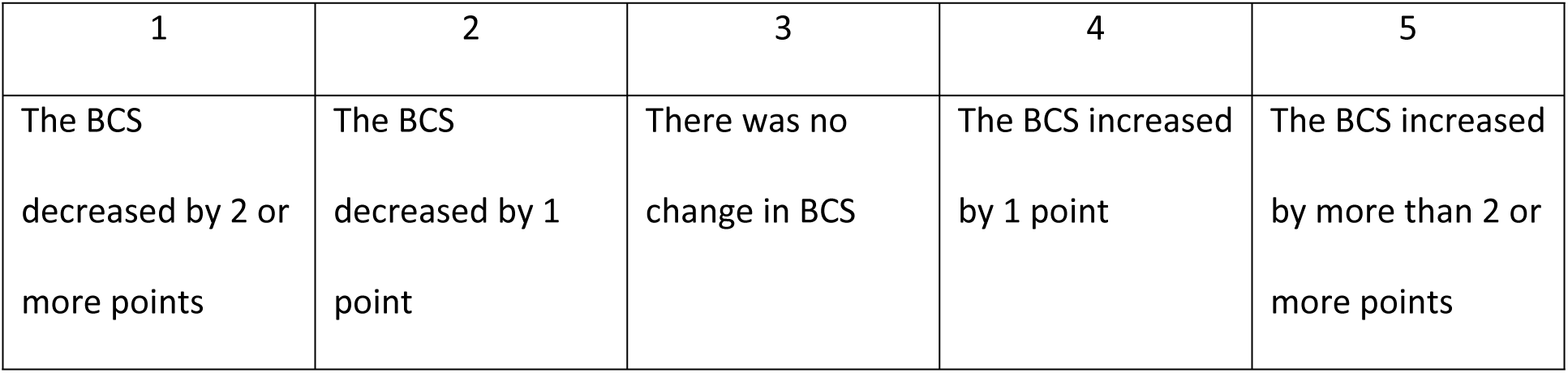

### Activity level

Guardians were asked: How active was your dog before switching to the plant-based food? After switching to the plant-based food what changes in activity, if any, did you notice?

Options:

| 1 | 2 | 3 | 4 | 5 |
| --- | --- | --- | --- | --- |
| Activity<br>decreased a lot | Activity<br>decreased a bit | There was no<br>change my dog's<br>activity | Activity increased<br>a bit | Activity increased<br>a lot |

### Gastrointestinal signs

#### Defaecation frequency

Guardians were asked: How many times per day did your dog defaecate (pass motions) per day before you switched to the plant-based food?

After feeding the plant-based food what happened?

Options:

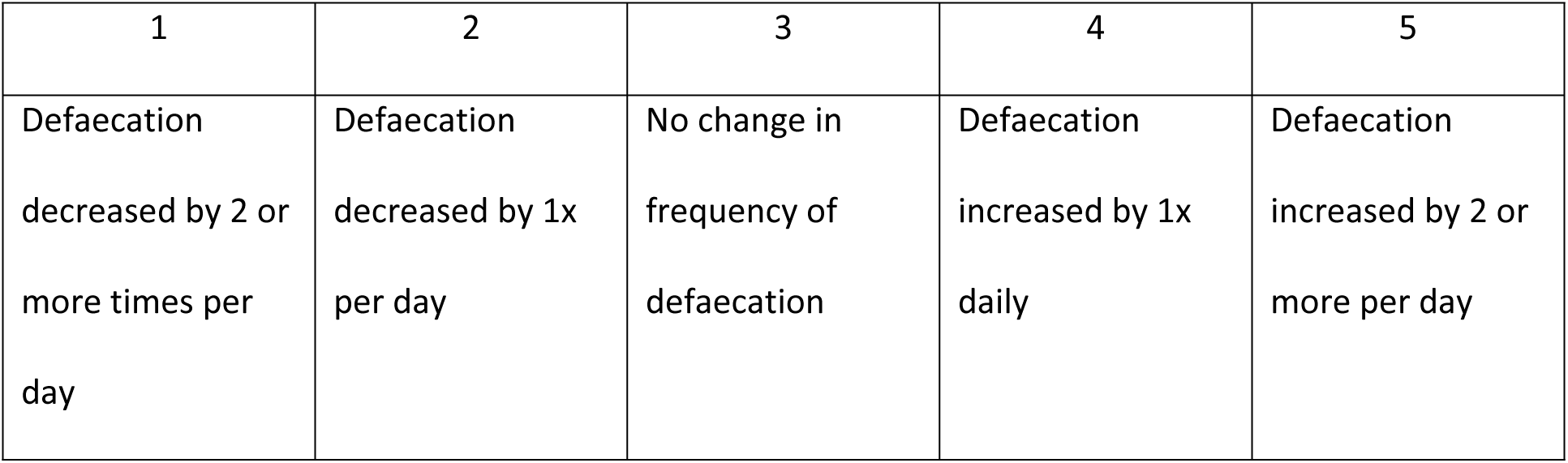

#### Faeces (stool) consistency

Guardians were asked: What was your dog’s faeces (stools) like before you switched to the plant-based food?

After switching to the plant-based food how did the consistency of faeces (stools) change?.

Options:

| 1 | 2 | 3 | 4 | 5 |
| --- | --- | --- | --- | --- |
| Faeces (stools)<br>became very<br>watery and loose<br>(diarrhoea) | Faeces (stools)<br>became softer<br>but were still<br>formed | No change in<br>consistency | Faeces (stools)<br>became firmer<br>and more well<br>formed | Faeces(stools)<br>became very hard<br>and dry |

#### Faecal colour

Guardians where asked: What colour were your dog’s faeces (stools) before you switched to the plant-based ration?

What happened to the faeces colour after you changed to the plant-based food?

#### Flatus (passing wind) frequency

Guardians were asked: Did your dog pass a lot of flatus “wind” before you switched to the plant-based food?

After switching to the plant-based food did you notice any change in frequency of flatus?

Options:

| 1 | 2 | 3 | 4 | 5 |
| --- | --- | --- | --- | --- |
| Passing wind<br>increased a lot | Passing wind<br>increased a bit | There was no<br>change in passing<br>wind | Passing wind<br>decreased a bit | Passing wind<br>decreased a lot |

#### Antisocial smelling flatus (wind)

Guardians were asked: Did your dog pass antisocial smelling wind before switching to the plant-based diet?

After switching to the plant-based food did you notice a change in the smell of “wind” passed by your dog?

Options:

| 1 | 2 | 3 | 4 | 5 |
| --- | --- | --- | --- | --- |
| The smell of wind got much worse | The smell of wind got a bit worse | There was no change in smell of wind passed by my dog | The smell of wind improved a bit | The smell of wind passed improved greatly |

### Dermatological signs

#### Haircoat Shine (Glossiness)

Guardians were asked: How shiny was your dog’s coat before switching to the plant-based food e.g dull, normal or shiny?

How did your dog’s haircoat change (if at all) following the change to plant-based food?

Options:

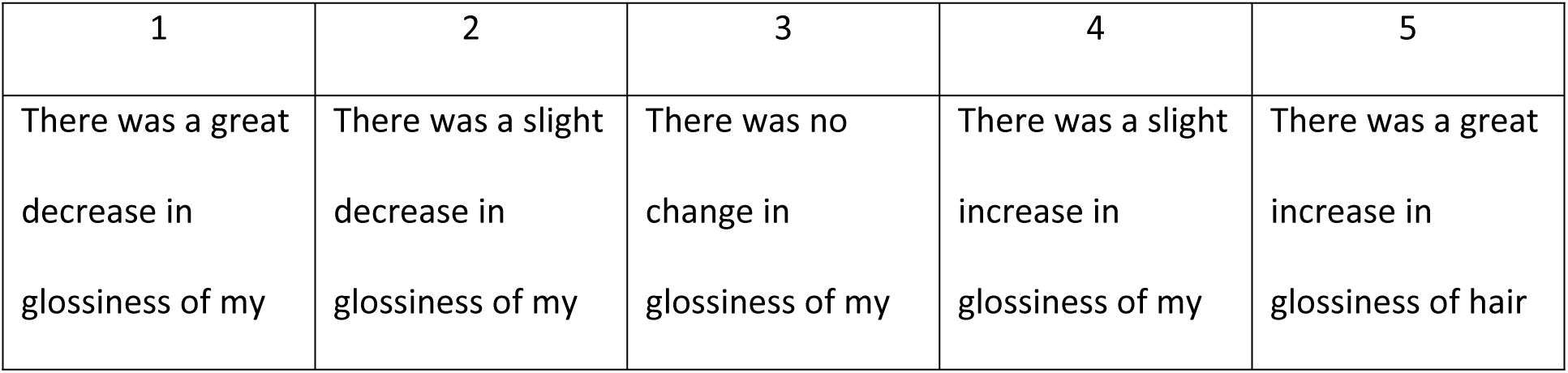
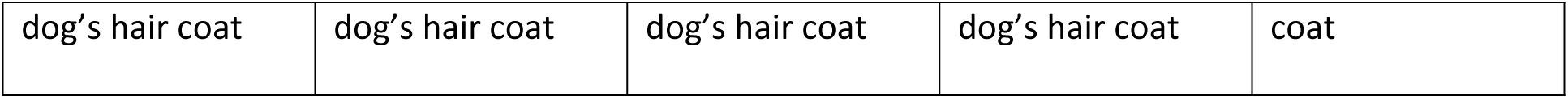

| 1 | 2 | 3 | 4 | 5 |
| --- | --- | --- | --- | --- |
| There was a great decrease in glossiness of my | There was a slight decrease in glossiness of my | There was no change in glossiness of my | There was a slight increase in glossiness of my | There was a great increase in glossiness of hair |
| dog's hair coat | dog's hair coat | dog's hair coat | dog's hair coat | coat |

#### Scale (Dandruff)

Guardians were asked: Did your dog have scales (dandruff) before switching to the plant-based food?

What happened afterwards?

Options:

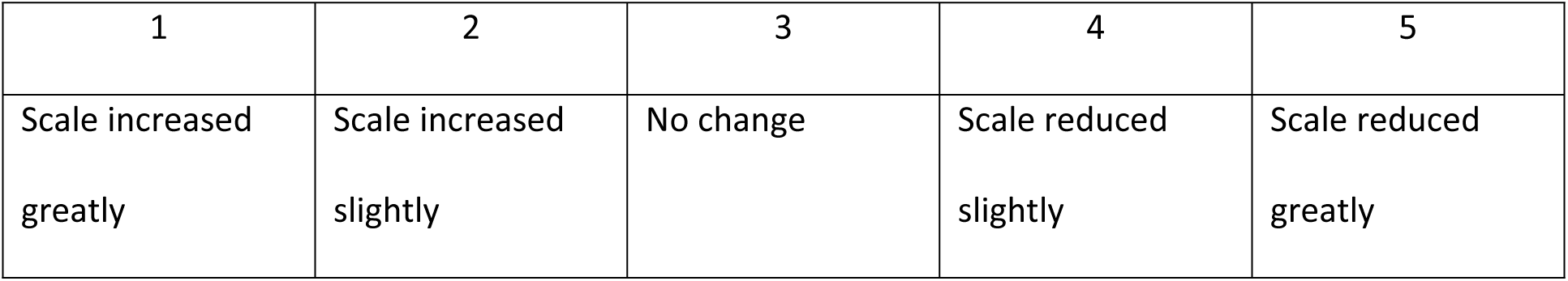

| 1 | 2 | 3 | 4 | 5 |
| --- | --- | --- | --- | --- |
| Scale increased<br>greatly | Scale increased<br>slightly | No change | Scale reduced<br>slightly | Scale reduced<br>greatly |

#### Crusting in ears (Otitis externa)

Guardians were asked did your dog have crusting, wax build up or redness in its ear canal before starting the plant-based food?

After switching to the plant-based food what changes if any did you notice

Options:

| 1 | 2 | 3 | 4 | 5 |
| --- | --- | --- | --- | --- |
| The amount of<br>crusting increased<br>a lot | The amount of<br>crusting increased<br>slightly | There was no<br>change in crusting<br>in the ear canal | The amount of<br>crusting reduced | The amount of<br>crusting resolved<br>totally |

#### Itchiness (Pruritus)

Guardians were asked whether their dog was itchy before switching to the plant-based food, and if so to report what change (if any) they noticed after switching

Options:

| 1 | 2 | 3 | 4 | 5 |
| --- | --- | --- | --- | --- |
| There was a great increase in itching | There was a slight increase in itching | There was no change in itching | There was a slight decrease in itching | The itching resolved totally |

#### Skin redness (erythema, inflammation)

Guardians were asked if their dog had skin redness (inflammation) before switching to the plant-based diet and if so what change (if any) did they notice:

Options:

| 1 | 2 | 3 | 4 | 5 |
| --- | --- | --- | --- | --- |
| A great increase in redness | A slight increase in redness | No change in redness | A slight reduction in redness | Total resolution of the redness |

### Behavioural signs

#### Aggression

Guardians were asked: After switching to the plant-based food have you noticed any changes in aggressive behaviour. If your dog never shows aggression you do not need to answer this question, please select ’Not relevant’ and move on

Options:

| 1 | 2 | 3 | 4 | 5 |
| --- | --- | --- | --- | --- |
| Aggressive<br>behaviour got a<br>lot worse | Aggressive<br>behaviour got a<br>bit worse | There was no<br>change aggressive<br>behaviour | The amount of<br>aggression<br>reduced | Previous<br>aggressive<br>behaviour<br>resolved |

#### Anxiety

Guardians were asked did your dog have signs of anxiety before starting plant-based food? If so, after switching to plant-based food what changes did you notice in anxiety?

Options:

| 1 | 2 | 3 | 4 | 5 |
| --- | --- | --- | --- | --- |
| The amount of<br>anxiety got much<br>worse | The amount of<br>anxiety got<br>slightly worse | There was no<br>change in anxiety | The amount of<br>anxiety reduced<br>slightly | My dog stopped<br>showing any signs<br>of anxiety |

#### Other Behaviour

Guardians were asked: Did your dog have other behaviour problems before starting the plant-based food e.g coprophagia (eating poo), chewing furniture? If so, please specify:

After switching to plant-based food what changes did you notice in behaviour?

Options:

| 1 | 2 | 3 | 4 | 5 |
| --- | --- | --- | --- | --- |
| The behavioural<br>problem got<br>much worse | The behavioural<br>problem got<br>slightly worse | There was no<br>change in<br>behaviour | The behavioural<br>problem<br>improved slightly | The behavioural<br>problem resolved<br>totally |

### Statistical analysis

Statistical analysis of Likert-type scale data is controversial (Sullivan and others 2013) and “the best way to display the distribution of responses is to use a bar chart” (St Andrews University 2022). So, the results of this study are expressed in simple descriptors and bar chart graphic format and no attempt is being made to calculate statistical significance using p values.

## Results

### The Likert Scale-style survey

101 dog guardians completed the survey however It emerged that one had not actually started feeding the plant-based food and so their answers had to be deleted from the analysis. Some guardians failed to answer some sections, or did not answer questions correctly. An example of an inappropriate answer was when some respondents claimed that their dog was not itchy before switching diets, but then reported that there had been a great improvement afterwards. Clearly such a response was not possible if the dog was not initially itchy so those results had to be excluded from the analysis.

The survey response rate was 32.6%. In expressing the results all percentages have been rounded to one decimal point.

### Key

n=number of valid responses, out of 100 possible.

% = percent of the number of valid responses, expressed to one decimal point

NAn – Not answered

NAa – Not appropriate answer

NR – Not relevant

### Acceptability (n=100)

82 dogs (82%) ate the novel plant-based food with enthusiasm when presented with it for the first time. 10 dogs (10%) ate some of the food when it was first presented then went back and finished it off later, 7 dogs (7%) ate the food when it was introduced gradually over several days, one dog would only accept the food if it was mixed with other foods. None of the dogs refused to eat the food. So, this plant=based food was highly acceptable (palatable).

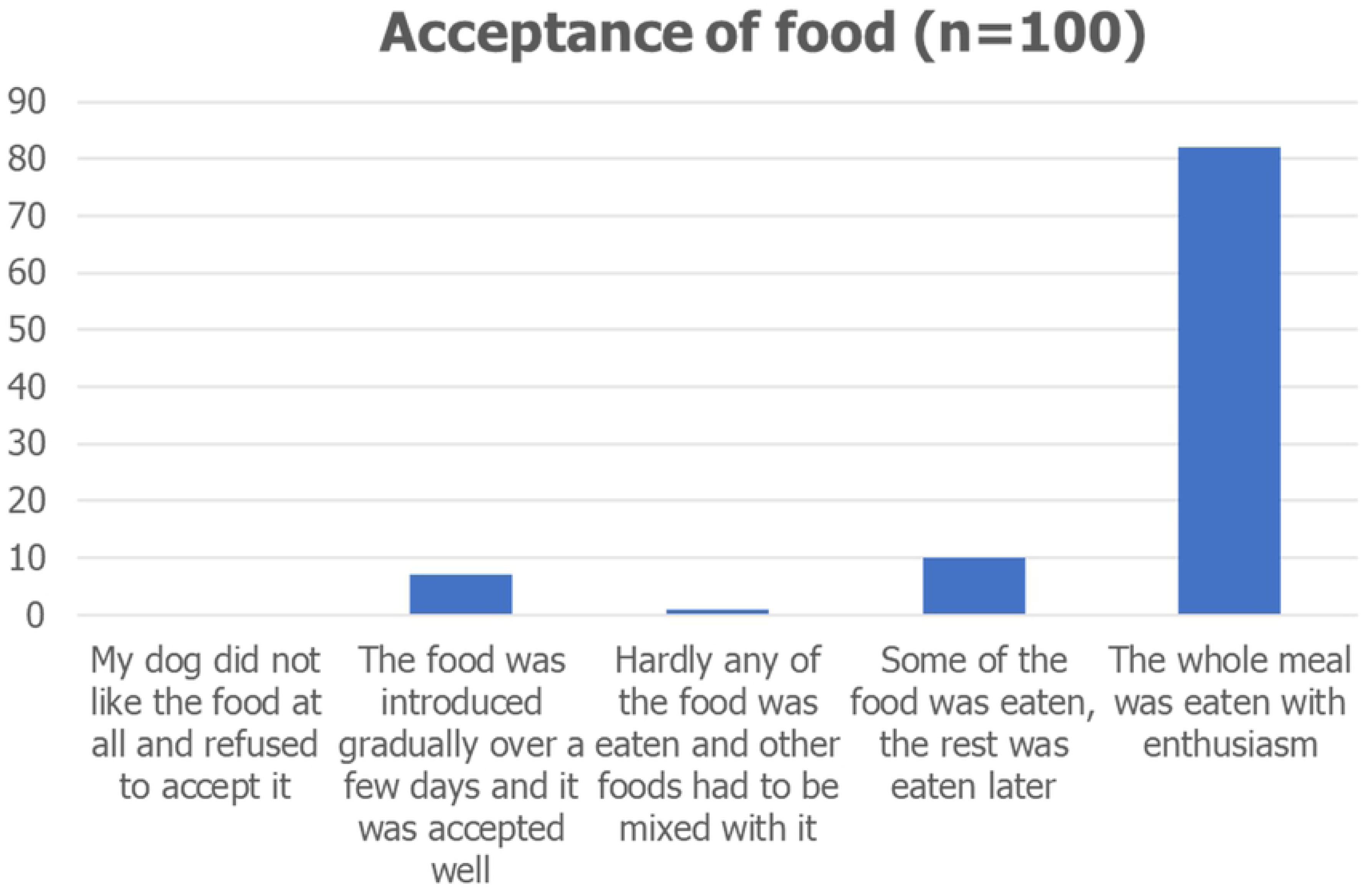

### Appetite (n=97) (NAn-3)

The majority (70 (72.2%)) of guardians reported that their dogs’ appetite did not change after switching to the plant-based food. 16 guardians (16.5%) reported a slight increase in their dogs’ appetite and 7 guardians (7.2%) reported a great increase in appetite, with their dog showing begging behaviour for more food. 4 guardians (4.1%) reported that their dog ate slightly less after switching. No dogs lost their appetite totally.

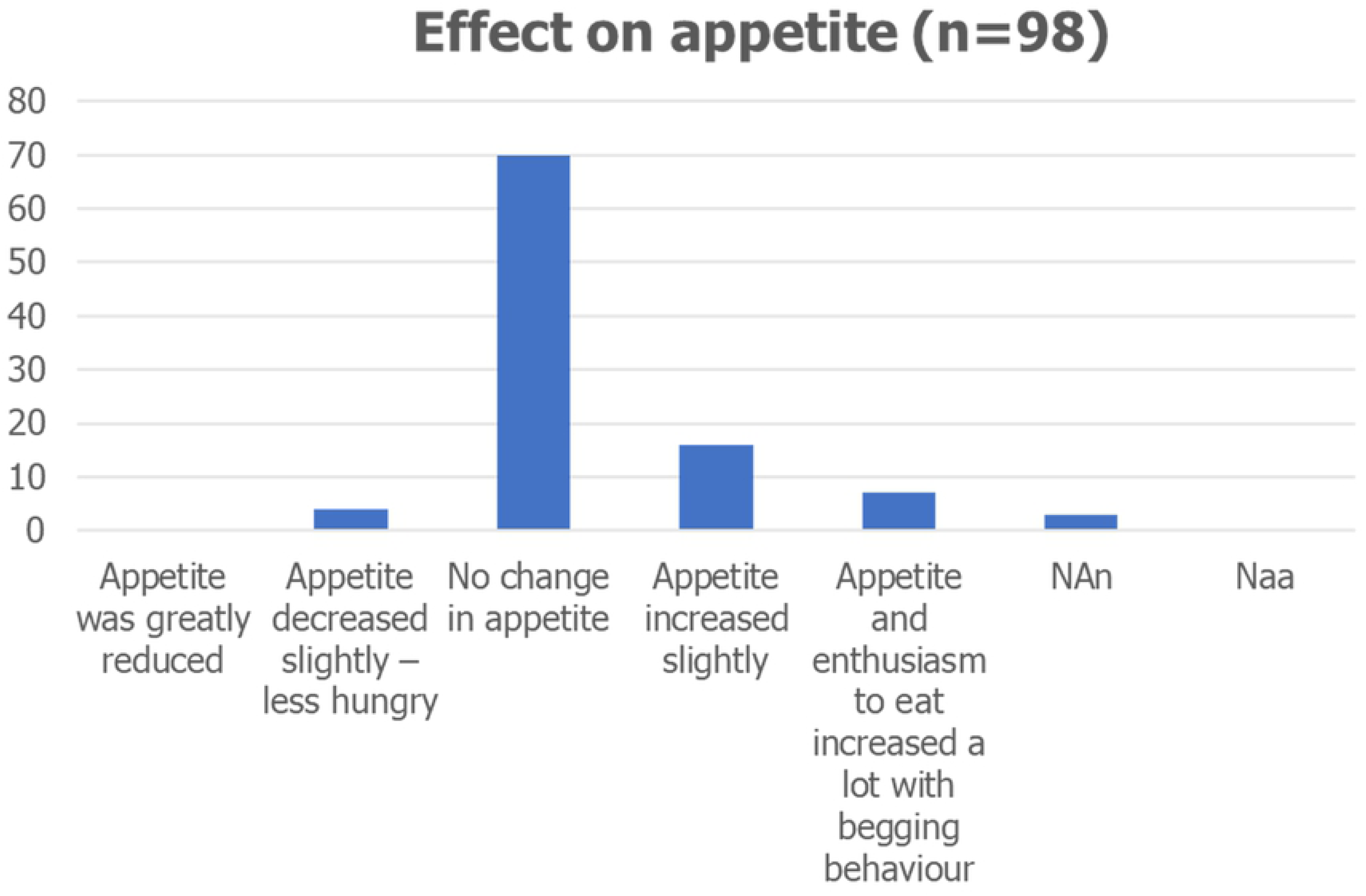

### Body Weight Changes (n=100)

71 guardians (71%) reported no change in body weight following a change to the plant-based food. 17 (17%) reported a drop in weight of which 4 (4%) dogs lost more than 10% of initial weight and 13 (13%) less than 10% weight. 12 guardians (12%) reported that their dogs had gained weight on the plant-based food, of which 8 (8%) gained less than 10% weight and 4 (4%) gained more than 10% of initial weight.

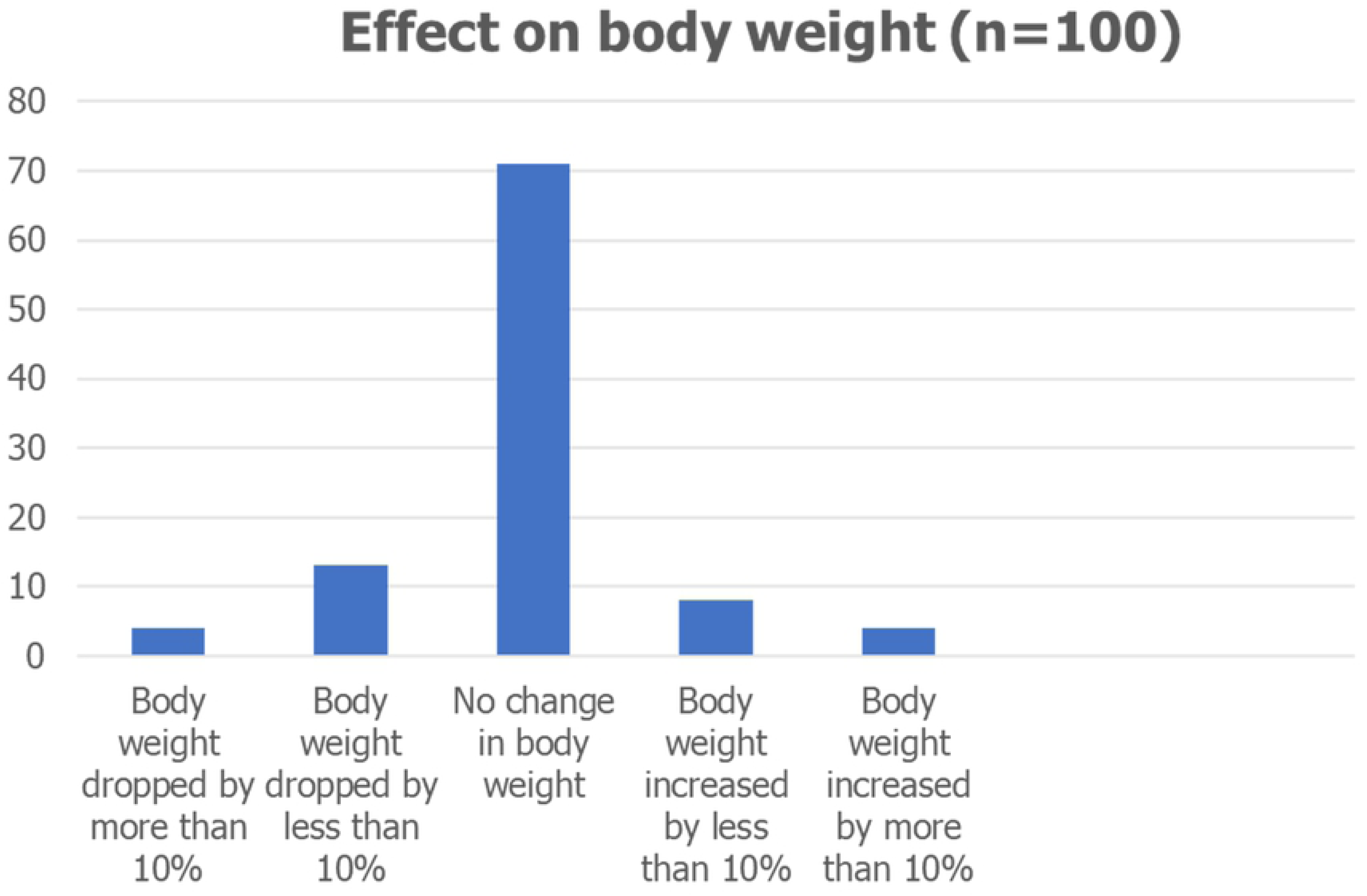

### Body Condition Score (BCS) (n=20)

20 guardians (20%) claimed they were using an accredited BCS score chart and completed this section. In 13 (65%) of these dogs BCS did not change, 3 dogs (15%) gained 2 or more BCS points, 3 (15%) gained one BCS point, and one dog (5%) lost 1 BCS point

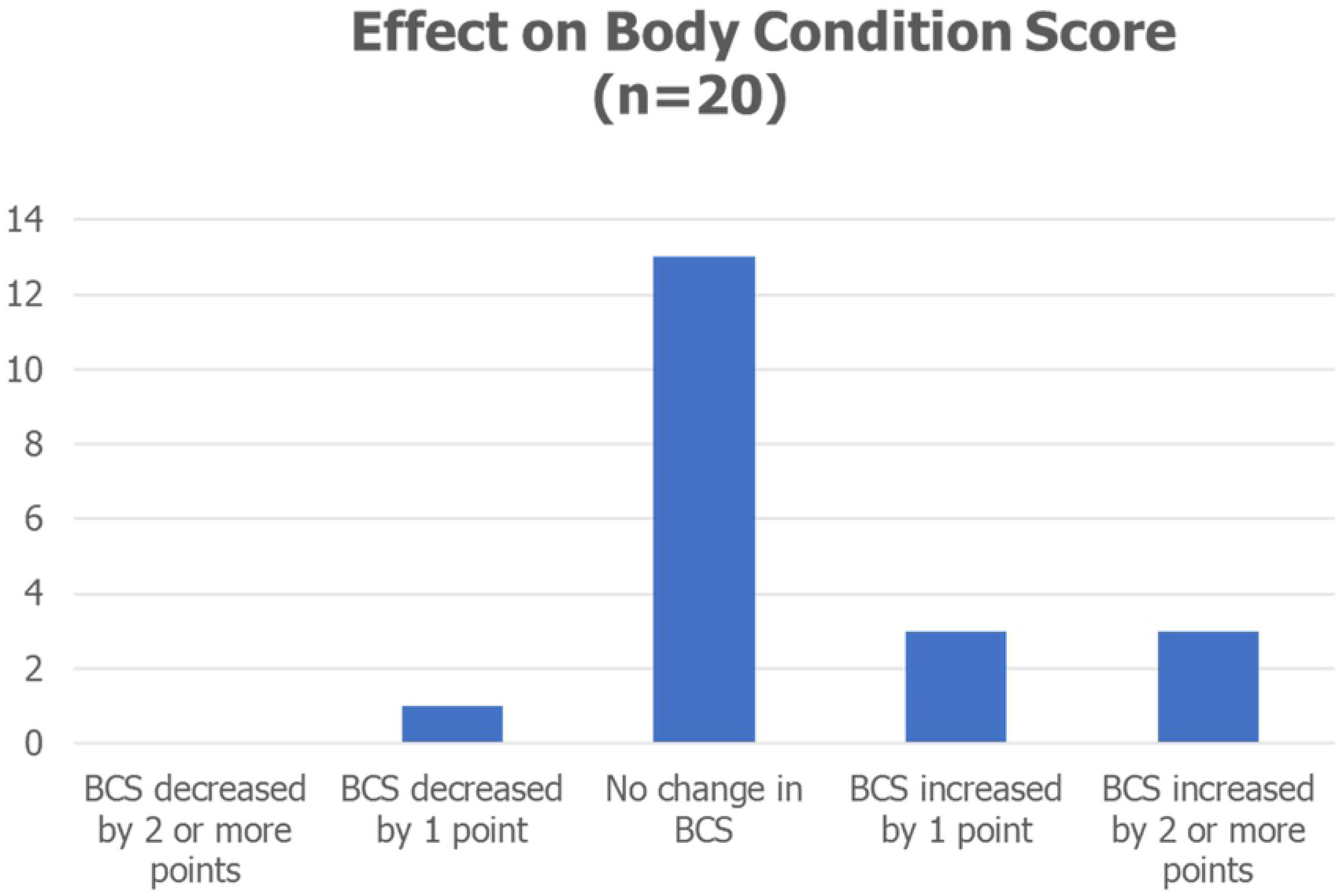

### Activity level (n=97) (NAn=3)

Following a switch to the plant-based food 67 (69.1%) dogs did not exhibit any change in activity. An increase in activity was reported for 28 (28.9%) dogs, in 19 (19.6%) activity was slightly increased and 9 (9.3%) became much more active. 4 (4.1%) of these dogs were not very active before the change in diet, 9 (9.3%) were active already and 3 (3.1%) were very active. Two dogs (2.1%) were reported to be slightly less active after the change in food. No dogs were reported to be a lot less active after switching to the vegan food.

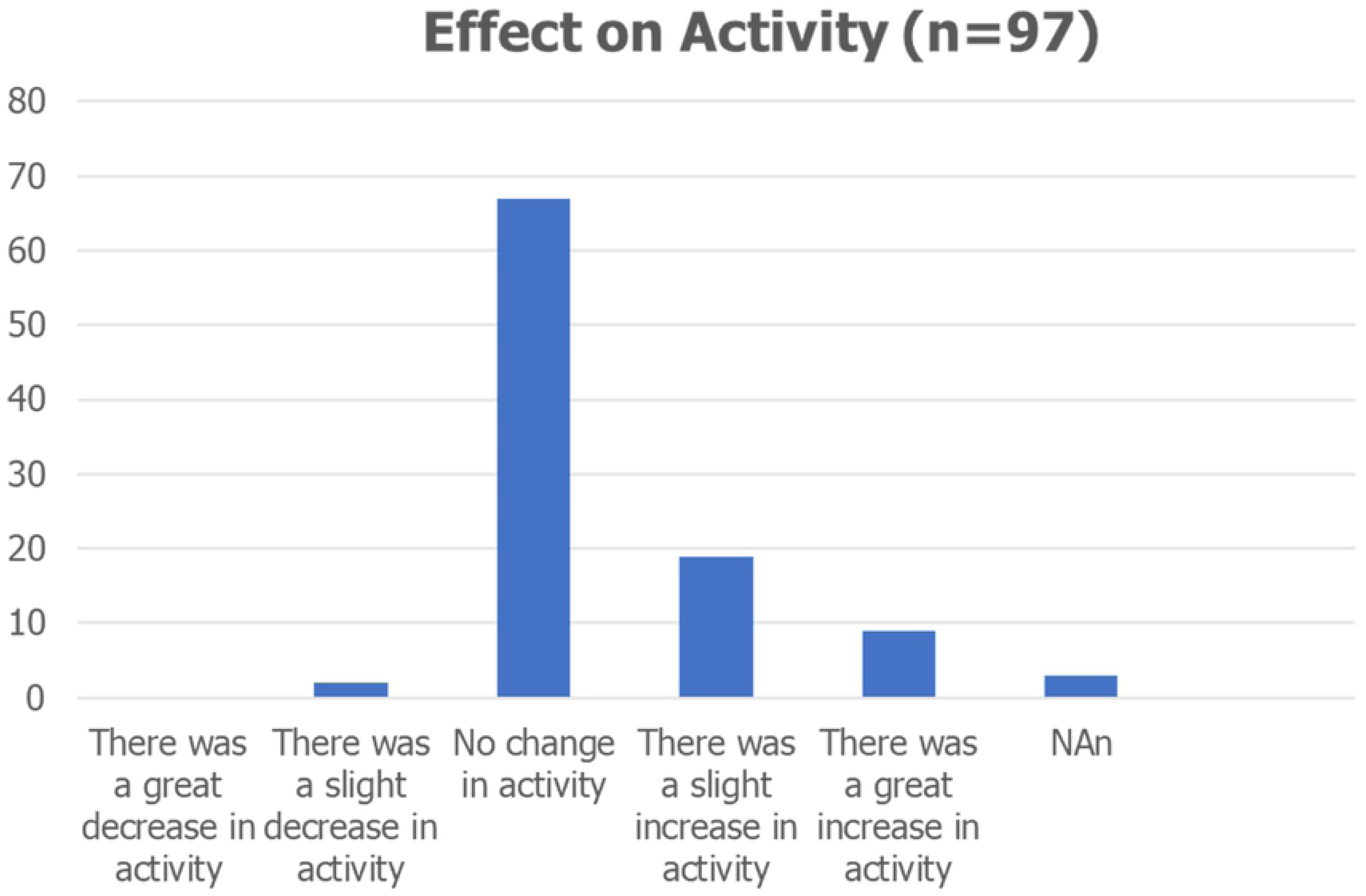

### Gastrointestinal signs

#### Defaecation frequency (n=94) (NAn=6)

Prior to switching diets 9 dogs (9.6%) were defaecating once per day, 32 (34.0%) twice per day, 33 (35.1%) three times per day, 12 (12.8%) four times per day, 3 (3.2%) five times per day, 4 (4.3%) six times per day and 1 (1.1%) eight times per day.

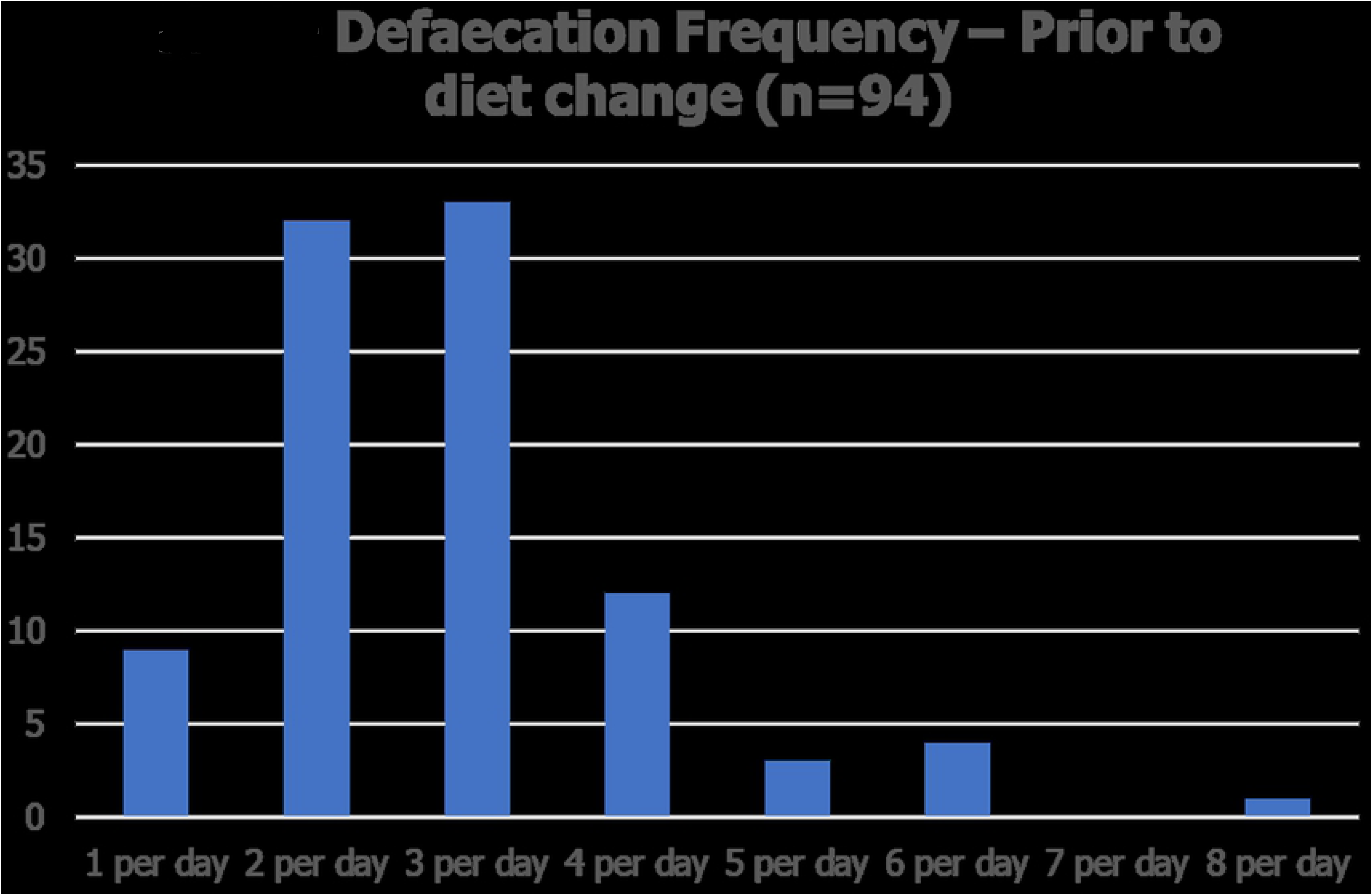
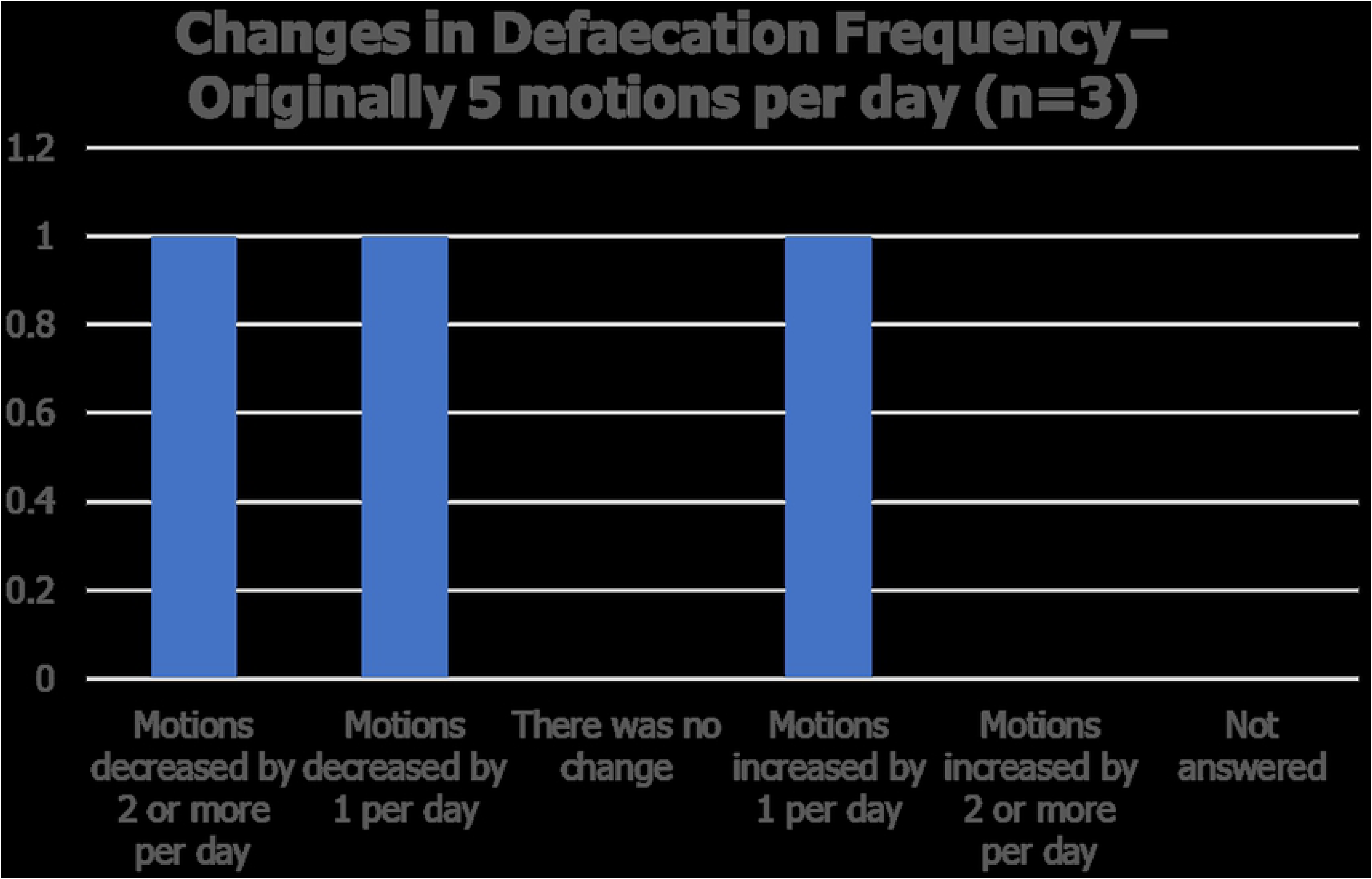
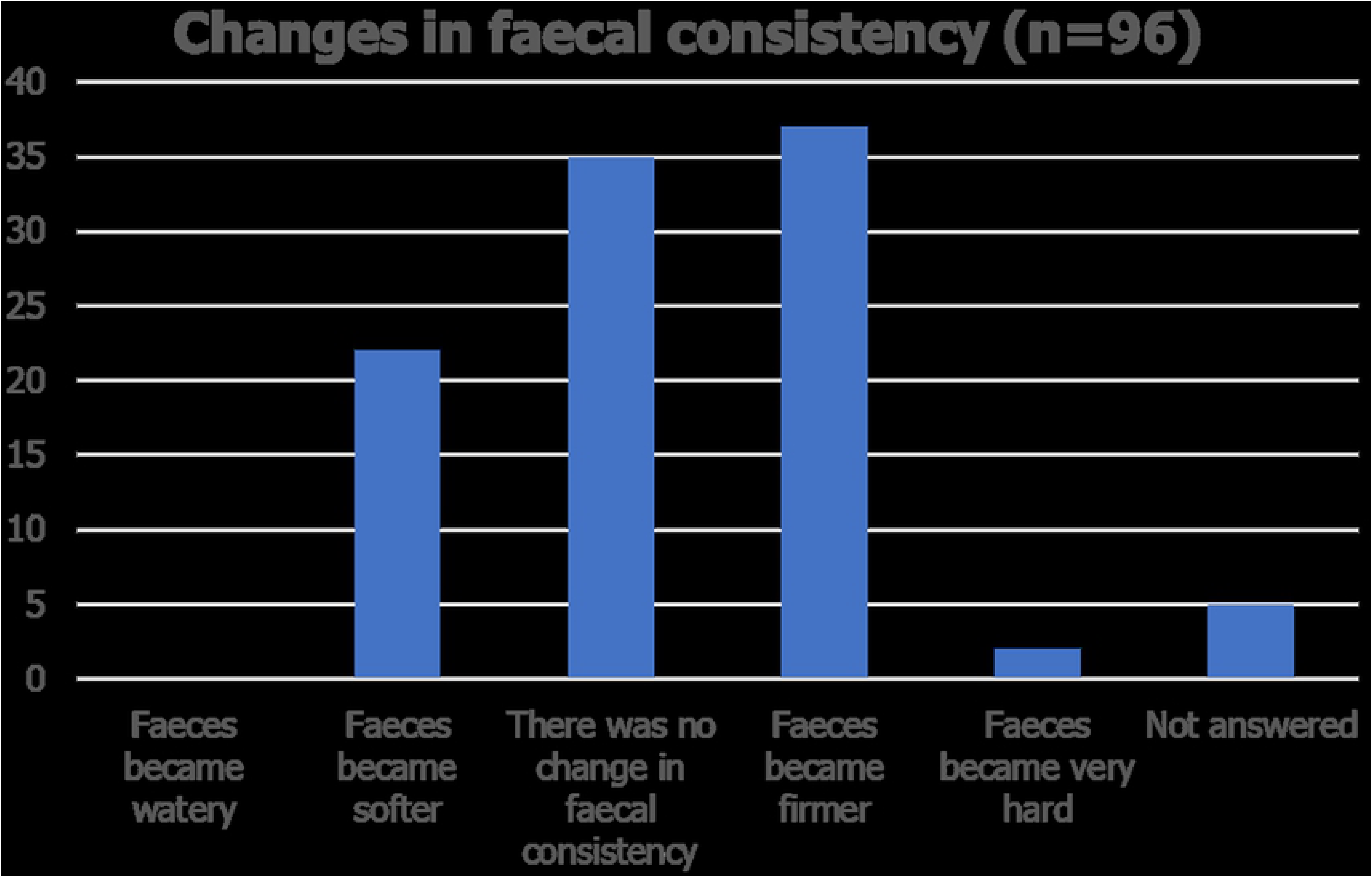

Of the 94 guardians who completed this section 53 (56.4%) reported that there had been no change in defaecation frequency after switching to the plant-based food. 11 guardians (11.7%) reported that their dog passed one more motion per day, and 6 (6.4%) that their dog passed 2 or more additional motions per day. The number of motions passed per day decreased by 1 in 16 dogs (17.0%) and by 2 or more per day in 8 dogs (8.5%).

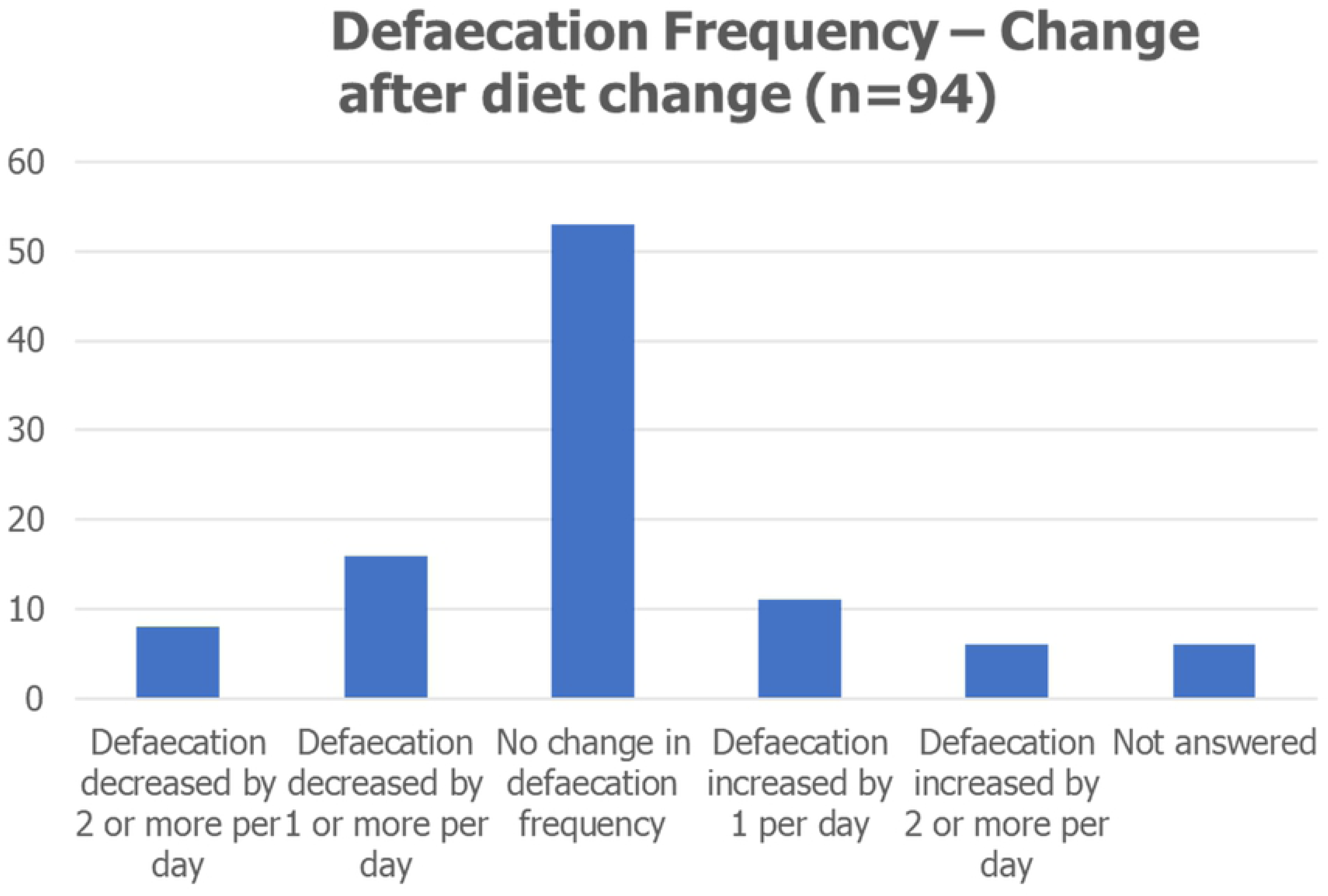
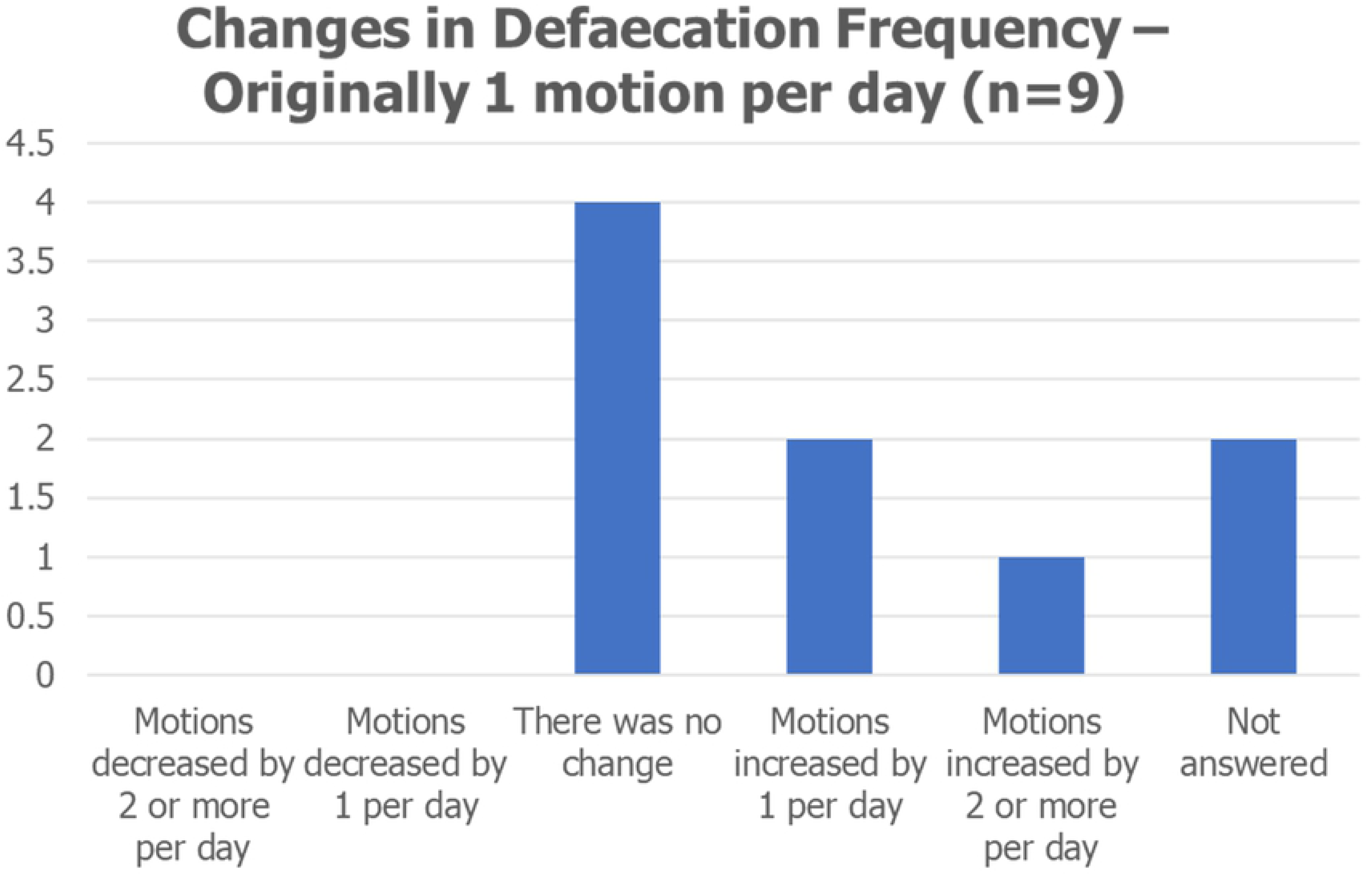
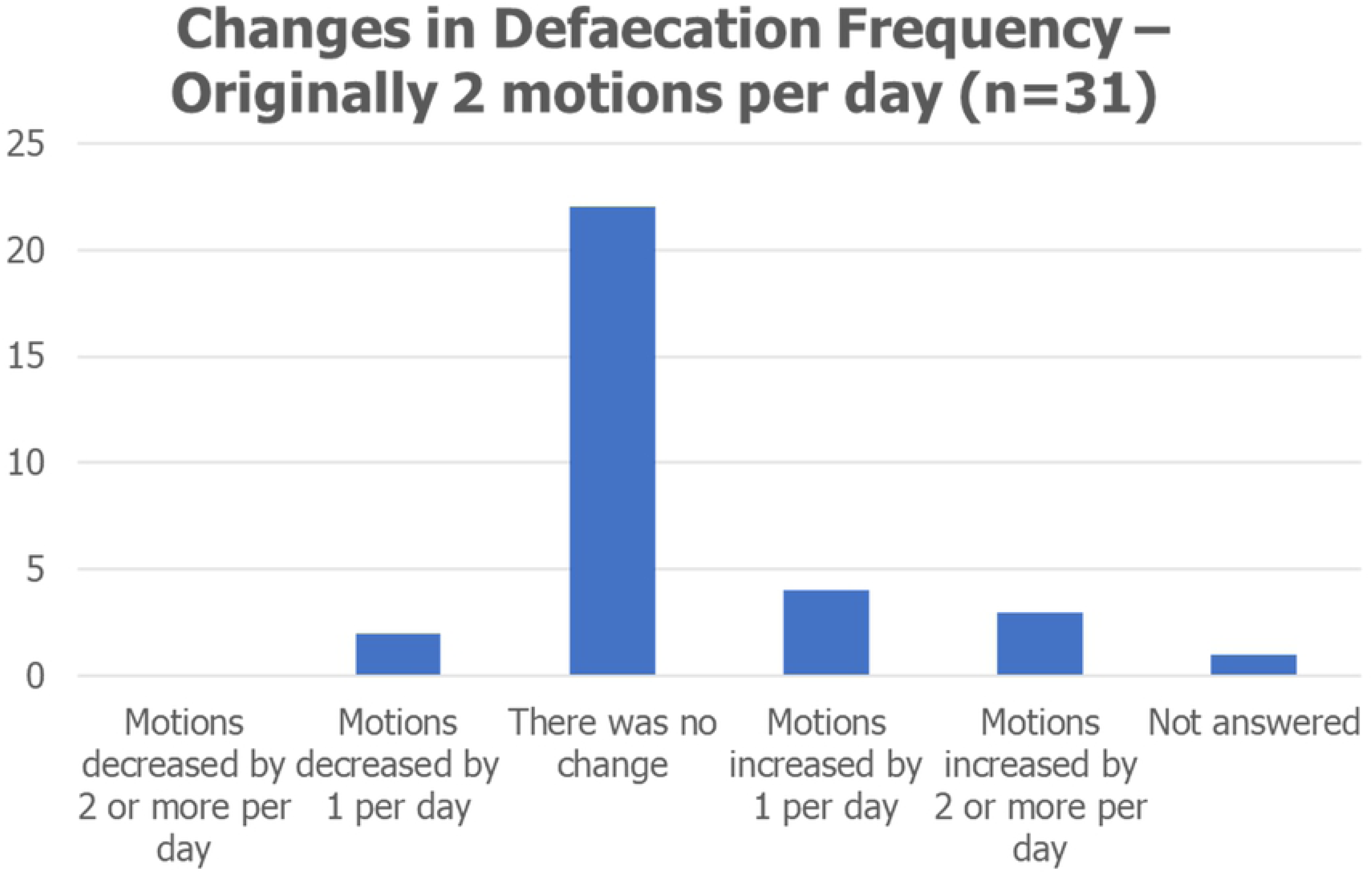
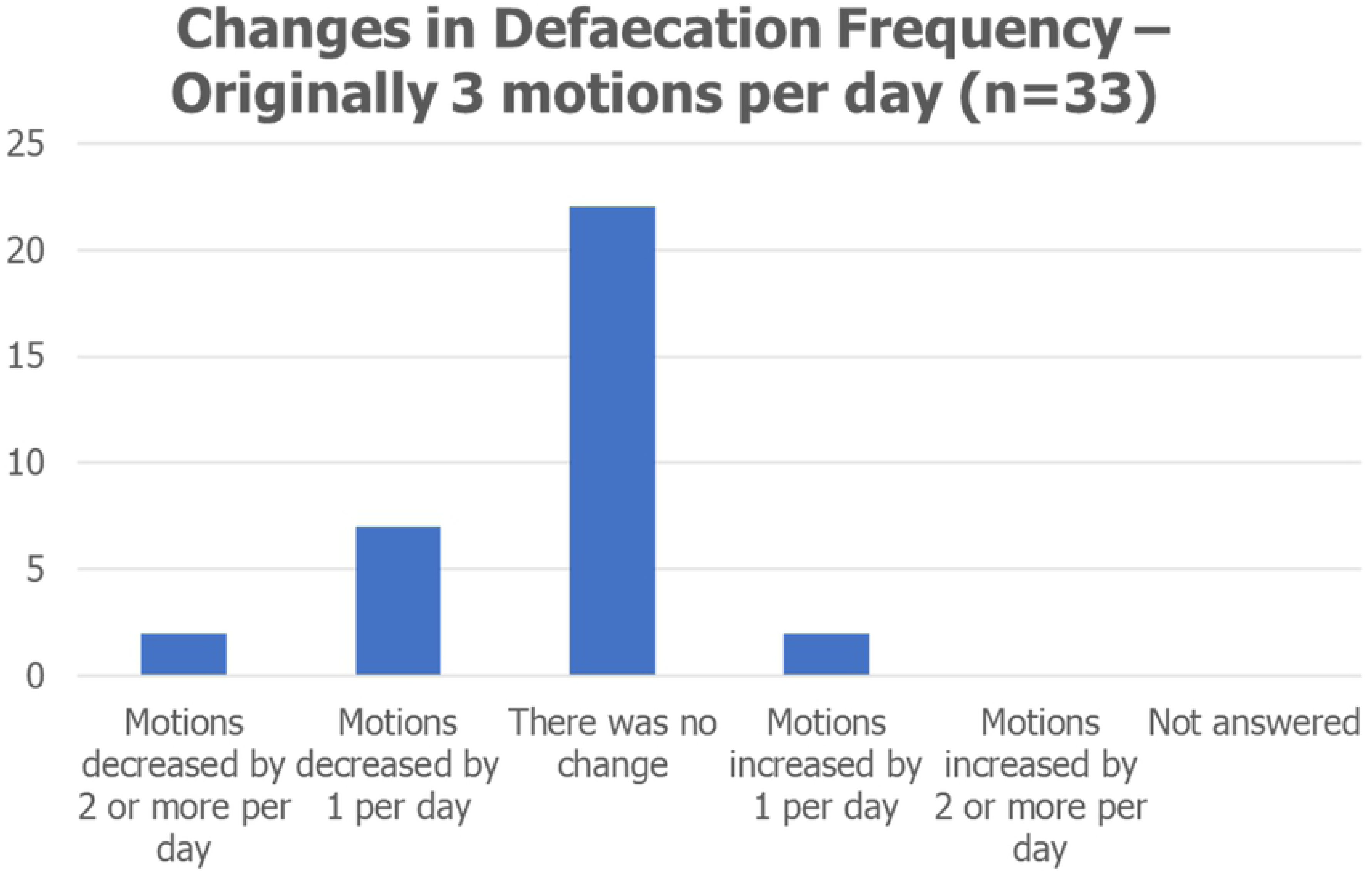
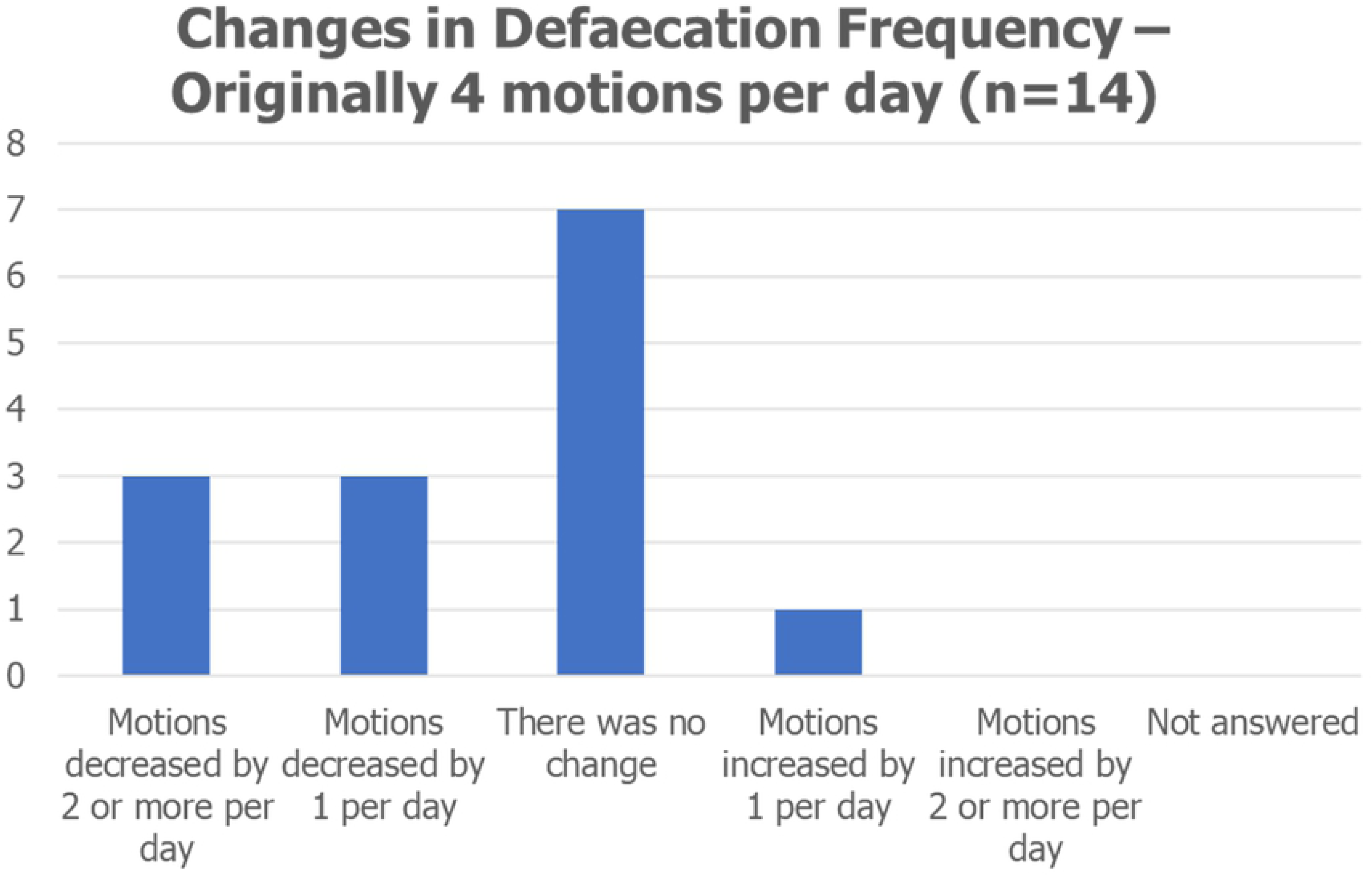
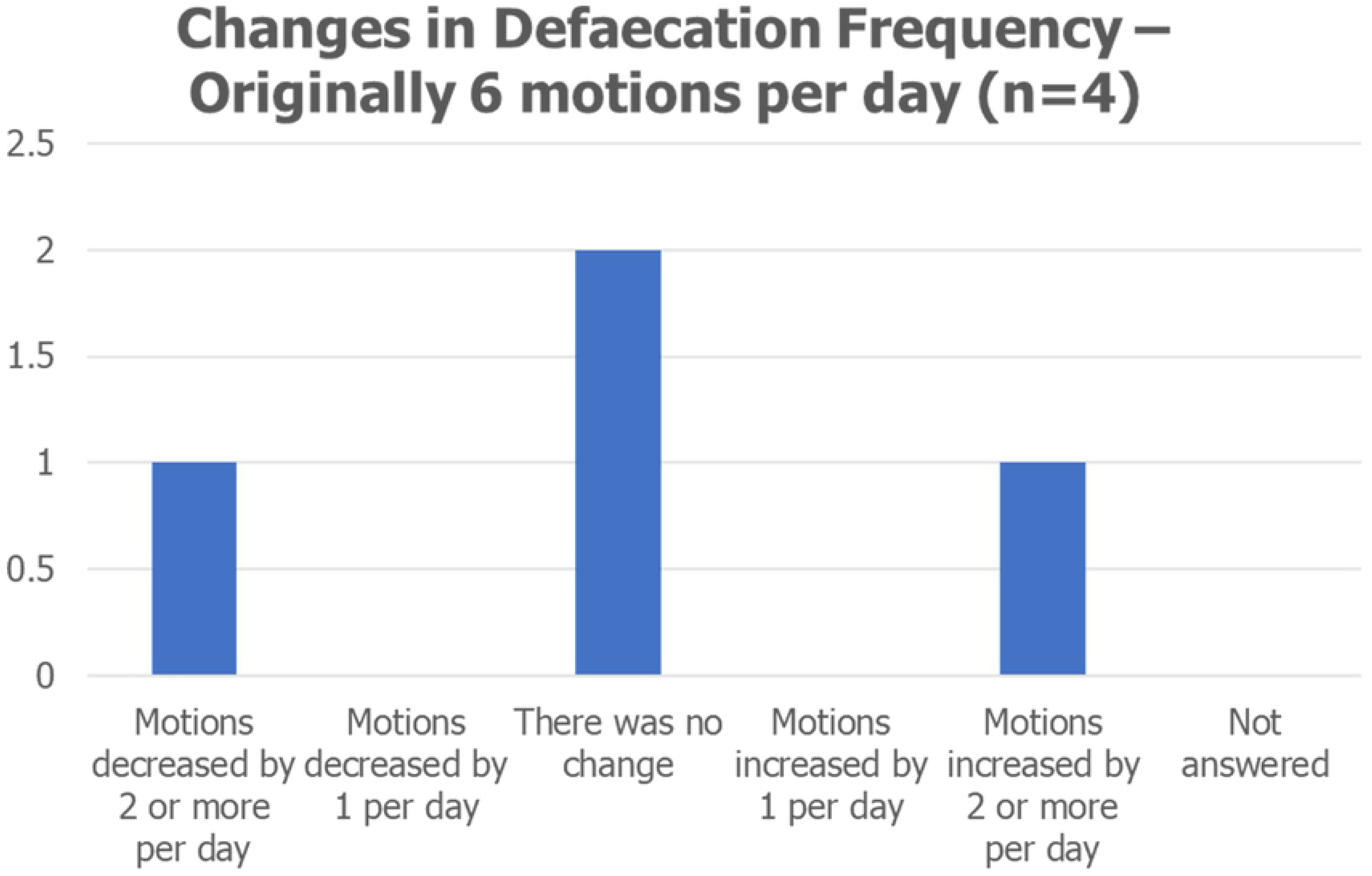
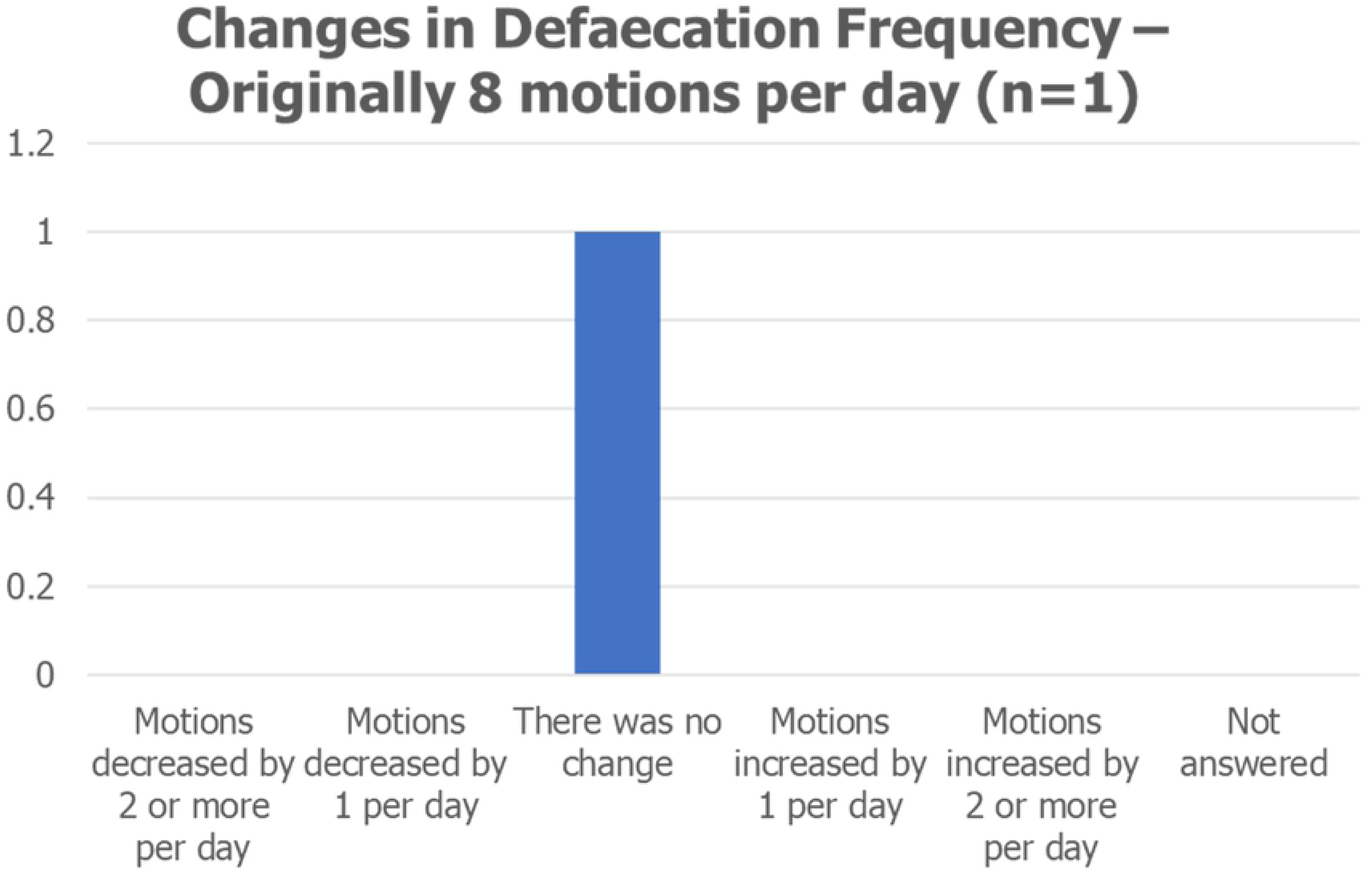

#### Faecal consistency (n=96) (NAn=4)

Prior to switching to a plant-based food 52 (54.2%) guardians described their dog’s faeces as being of normal consistency, 22 (22.9%) as soft, 10 (10.4%) as watery, 9 (9.4%) as hard and 2 (2.1%) as very hard. After switching a total of 35 guardians (36.5%) reported no change in faecal consistency, 37 (38.5%) reported the faeces was firmer, 2 (2.1%) that the faeces was much harder, 22 (22.9%) reported that the faeces was softer and no guardians reported that the faeces became watery.

In the dogs with normal faeces there was no change in consistency in 33 (63.5%), in 8 (15.2%) the stools became firmer and 11 (21.2%) softer after diet change.

In dogs with watery faeces before switching to plant-based ration there was no change in one dog but 9 out of ten (90.0%) developed firm faeces on the plant-based food. For dogs with soft faeces prior to diet change 19 out of 22 (86.4%) developed firm faeces after the change, there was no change in consistency in two (10.5%) of the dogs and in one dog (5.2%) faeces got softer.

For dogs with hard faeces initially 6 out of 9 (66.7%) the dog’s faeces was softer but still formed after switching, in 2 (22.2%) dogs the faeces was harder after switching and in one there was no change. No guardians reported watery faeces after switching to the plant-based diet.

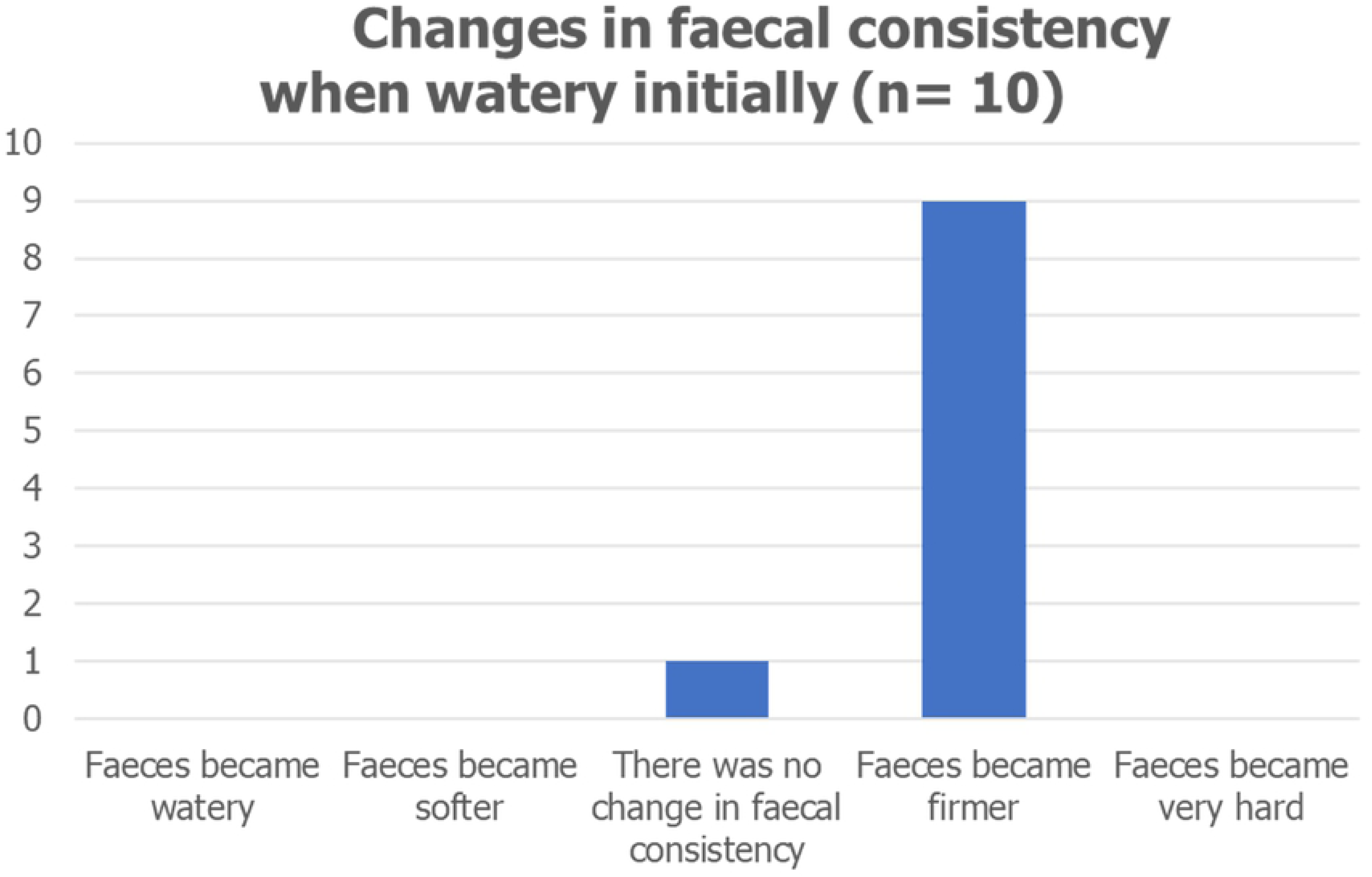
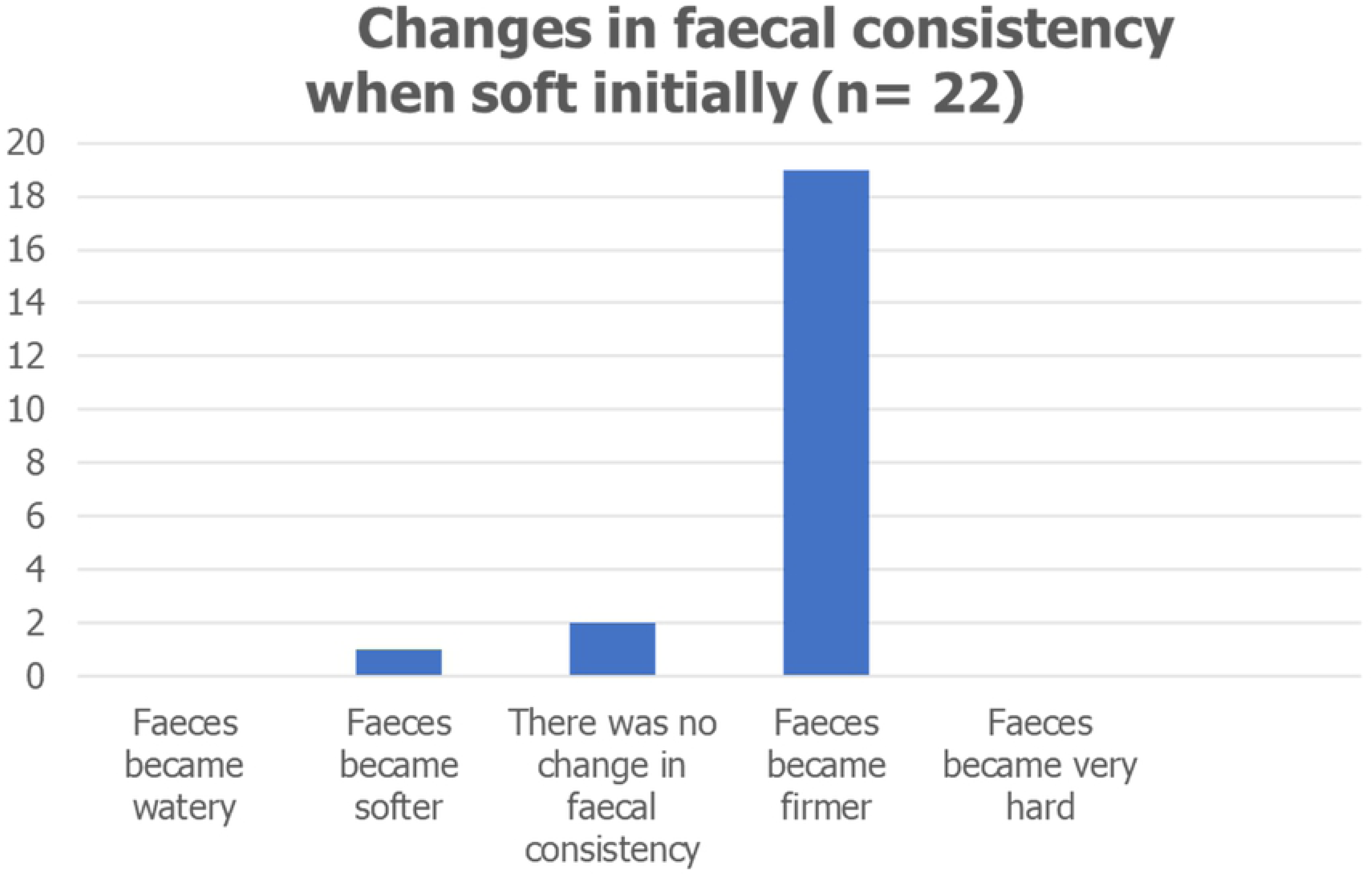
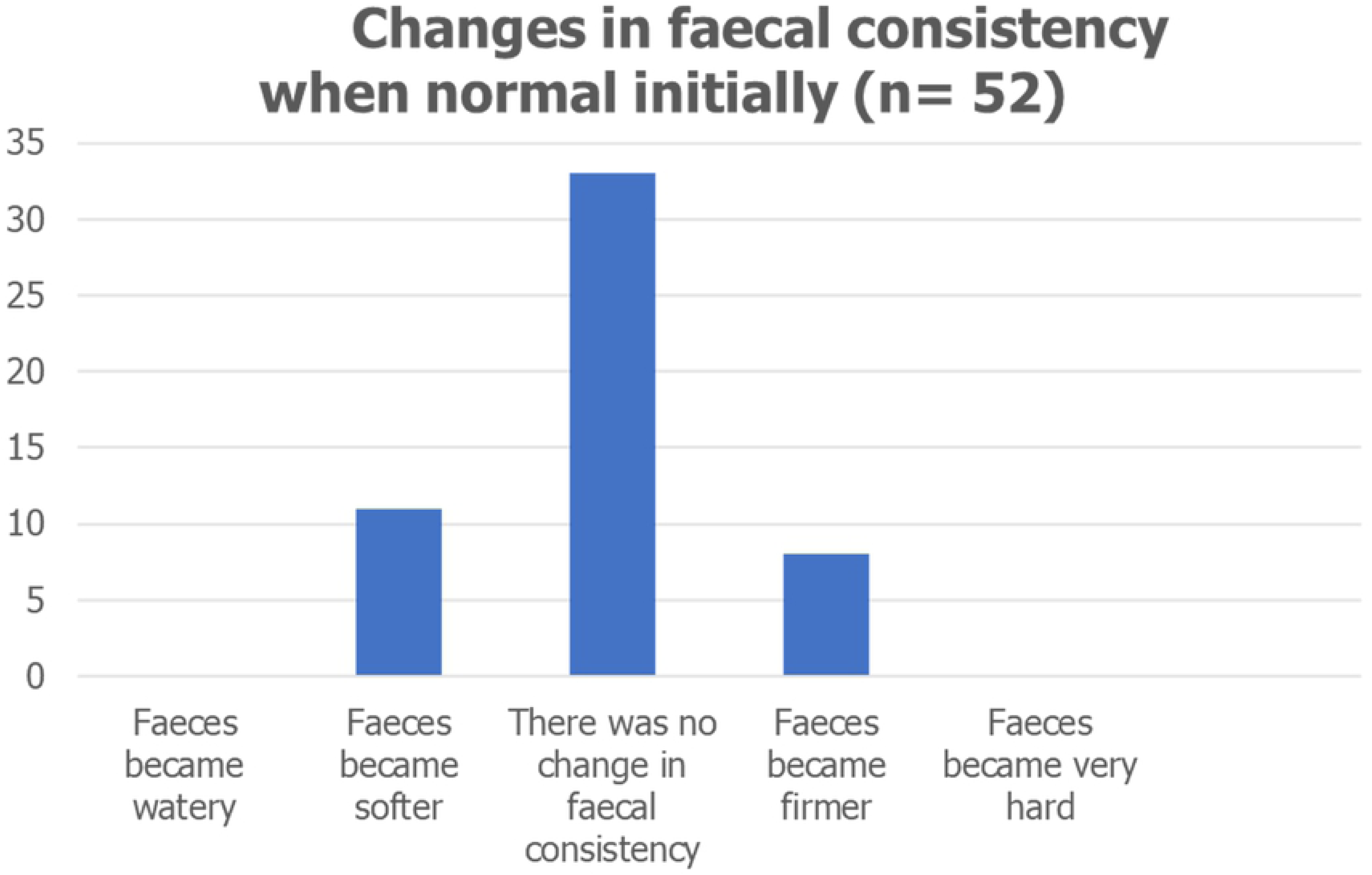
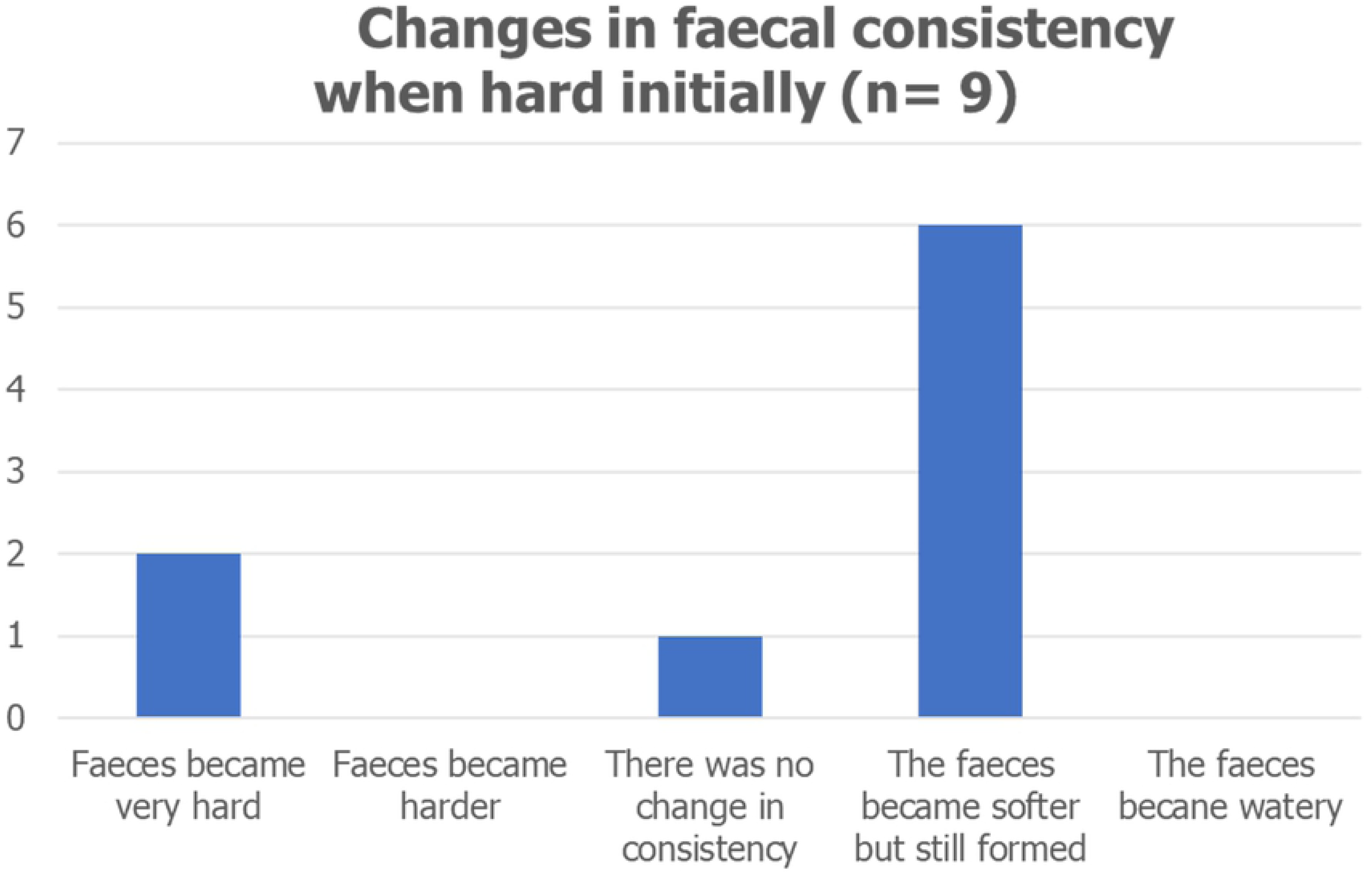

#### Faecal Colour (n=93) (NAn=7)

Prior to changing to the plant-based ration 47 (50.5%) of guardians described their dog’s faeces as being brown in colour, 18 (19.4%) described it as light brown and 18 (19.4%) described it as dark brown. 6 (6.5%) guardians claimed their dog had yellow faeces, 2 (2.2%) black faeces and 2 (2.2%) orange faeces.

Following a change to the plant-based food 39 guardians (41.9%) reported no change in colour, 29 (31.2%) said the faeces was lighter and 23 (24.7%) darker. Of the dogs with light brown faeces initially 15 (83.3%) got darker, and of the dogs with dark brown faeces 16 (88.9%) got lighter, of dogs with yellow faeces initially 3 (50.0%) changed to dark brown, 2 (33.3%) did not change, and one (16.7%) changed to light brown. In the two dogs with orange coloured faeces one did not change and the other became lighter in colour. Of the two dogs with black faeces in both the faeces became lighter.

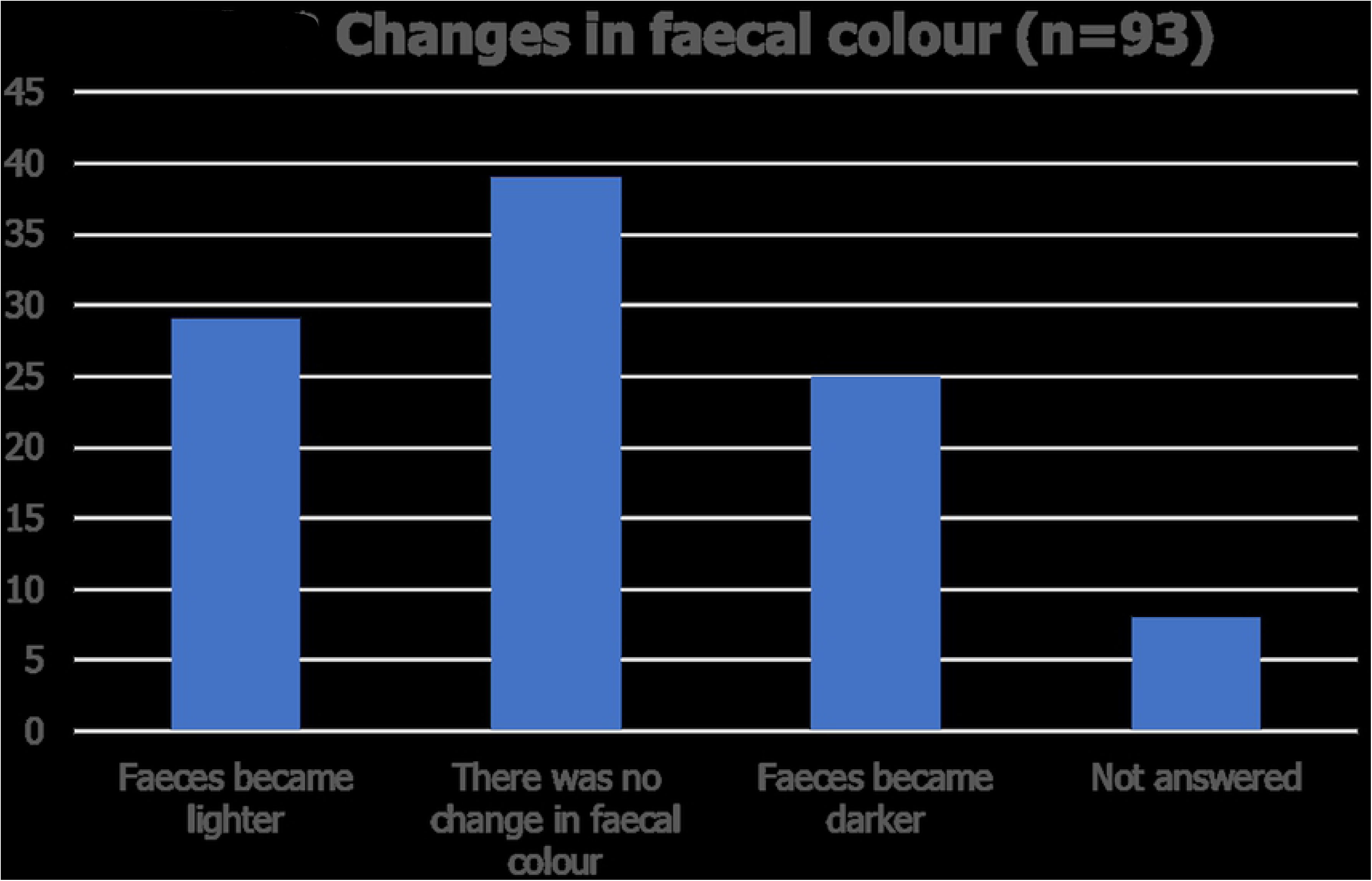

#### Flatus (passing wind) Frequency (n=96) (NAn=4)

Of the 96 dogs 74 (77.1% %) did not pass a lot of wind prior to diet change. Of these there was no change in 64 (86.5%), 8 (10.8%) passed more wind after the change in diet with 2 (2.7%) dogs passing a lot more.

22 (22.9%) guardians reported that their dogs passed a lot of wind prior to diet change and afterwards in these dog’s flatus was reduced a lot in 13 (59.1%), reduced slightly in a further 5 (22.7%) and there was no change in 4 (18.2%). None of these guardians reported an increase in passing wind after changing to a plant-based diet

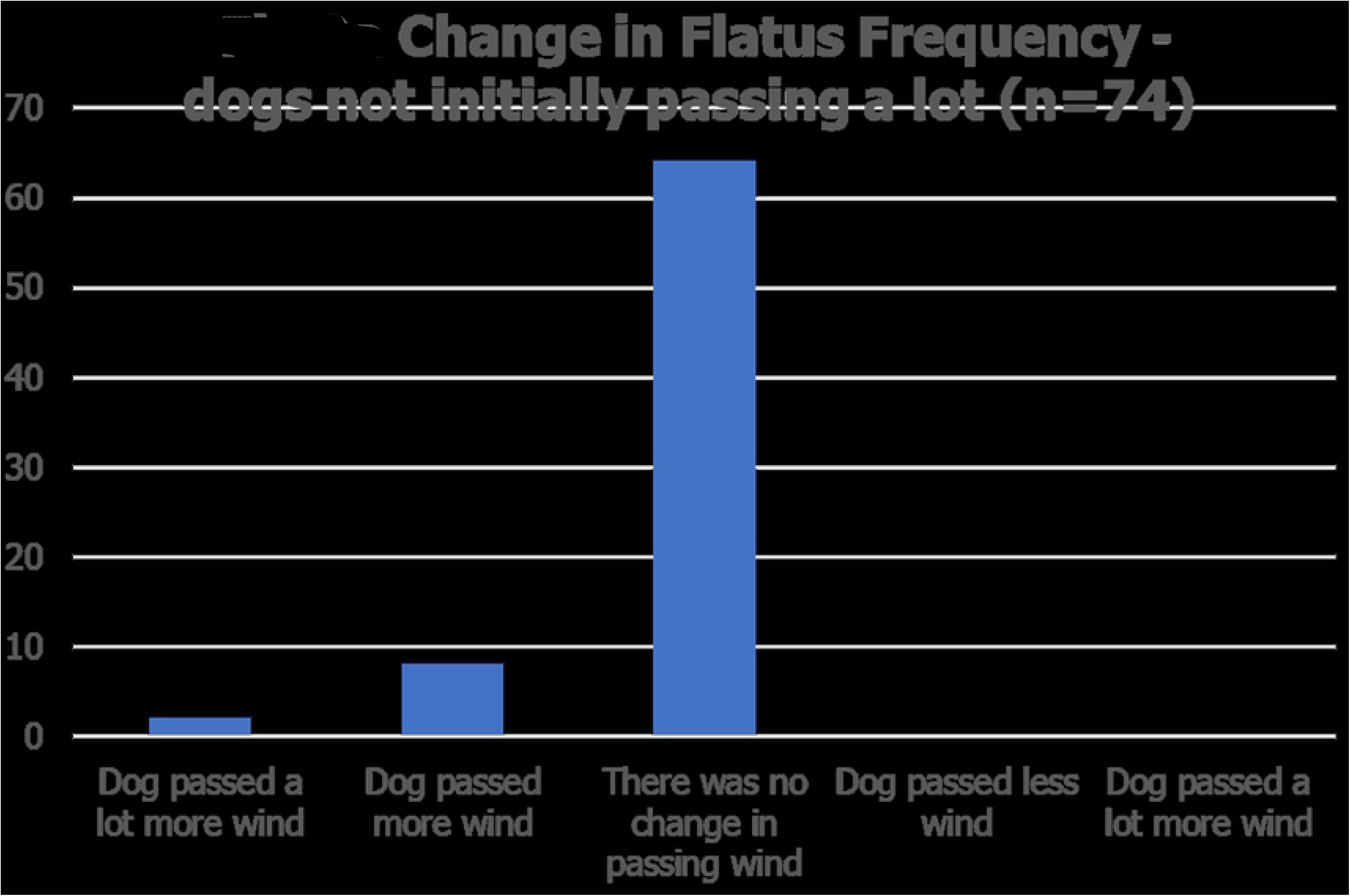
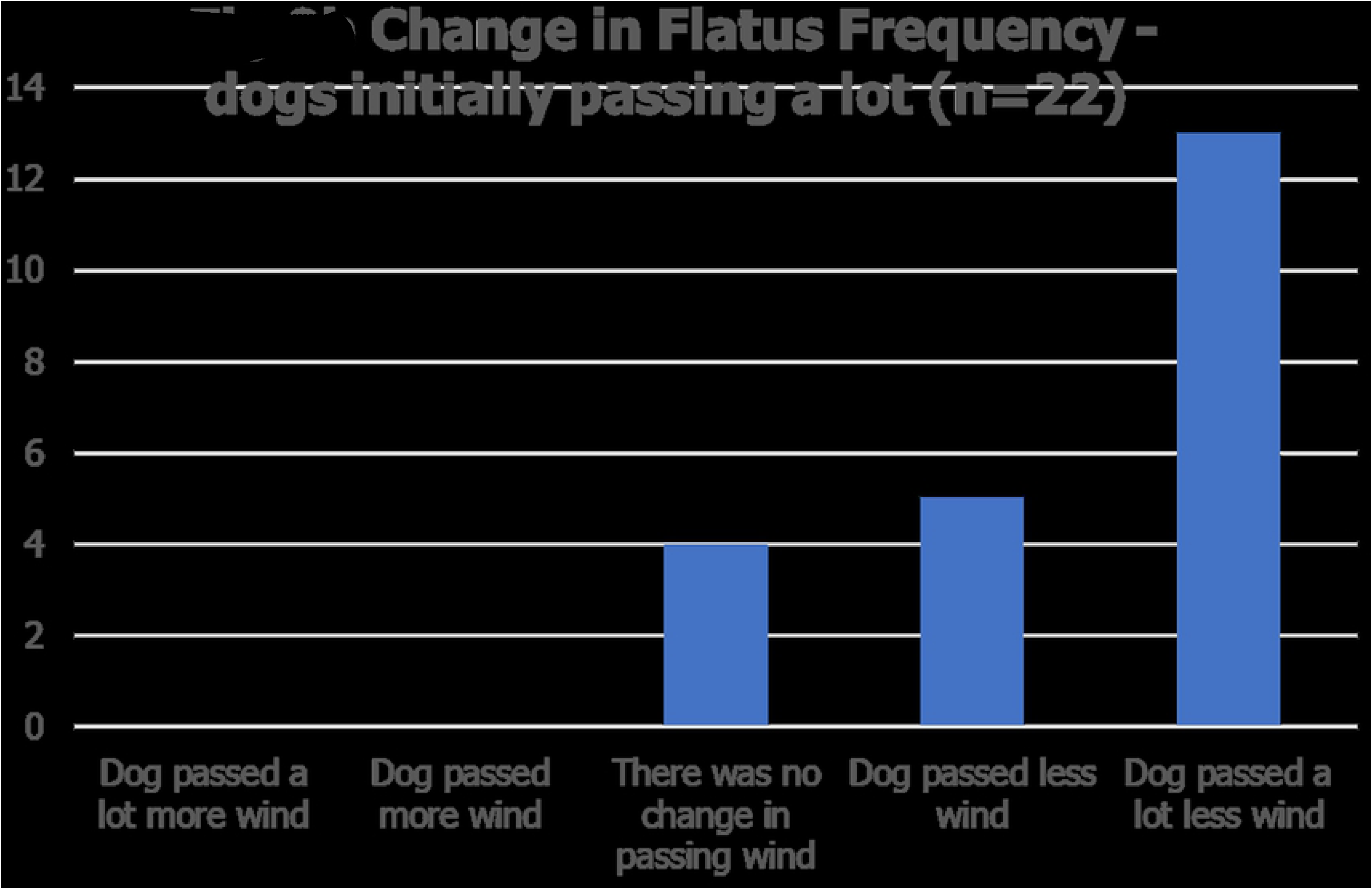

#### Antisocial smelling flatus (wind) (n=97) (NAn=3)

Of 97 guardians who completed this section 26 (26.8%) reported that their dog passed antisocial smelling wind before switching to a plant-based diet. After changing diet, in 19 (73.1%) of these the smell improved of which 17 (65.4%) reported that the smell improved greatly. In 7 (26.9%) of these dogs there was no change in smell. The smell did not get worse in any of the dogs.

71 (73.2%) of guardians reported that their dog did not pass antisocial smelling wind prior to a change to the plant-based food, afterwards there was no change in 65 (91.5%), the smell got slightly worse in 5 dogs (7.0%) and in one dog (1.4%) the smell improved a bit.

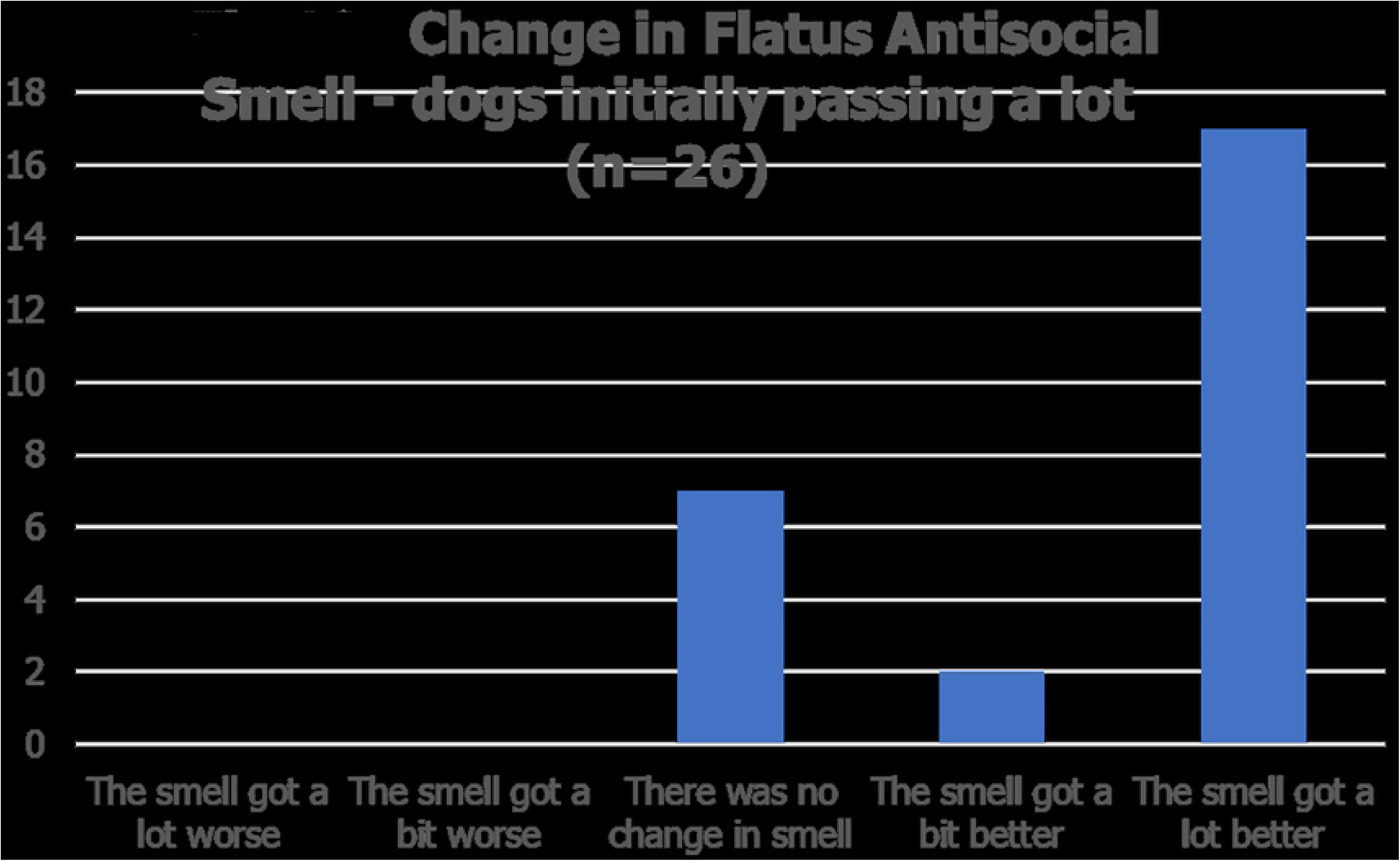
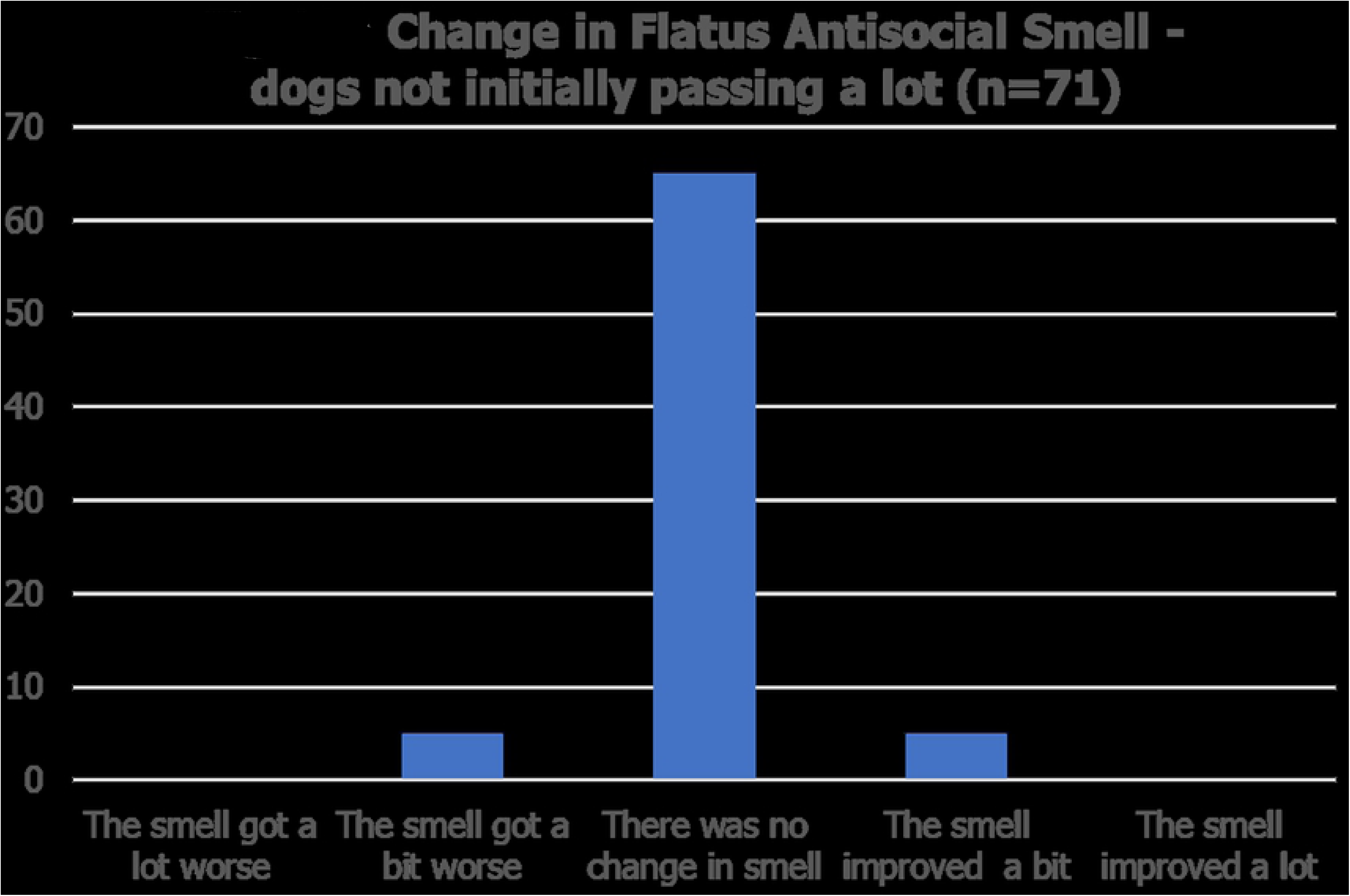

### Dermatological signs

#### Hair Coat Glossiness

##### a. Before switching to the plant-based food (n=83) (NAn=9; NAa= 8)

Prior to switching to the plant-based food 49 (59%) of guardians reported that their dog had normal hair shine, one (1.2%) said their dogs’ coat was very dull and 5 (6.0%) that the haircoat was dull, whereas 21 (25.3%) reported that their dog had shiny hair and 6 (7.2%) very shiny hair.

##### b. After switching to the plant-based food (n=98) (NAn=2)

After the switch to the plant-based ration 48 guardians (49.0%) reported an improvement in hair coat glossiness with 26 of these (26.5%) reporting a slight increase in shine and 22 (22.4%) reporting a great improvement in shine. Three guardians (3.1%) reported that their dogs’ hair coat appeared to be slightly duller after switching to the plant-based food. No guardians reported a great deterioration in hair shine. 47 guardians (48.0%) reported no change in hair shine after switching diets.

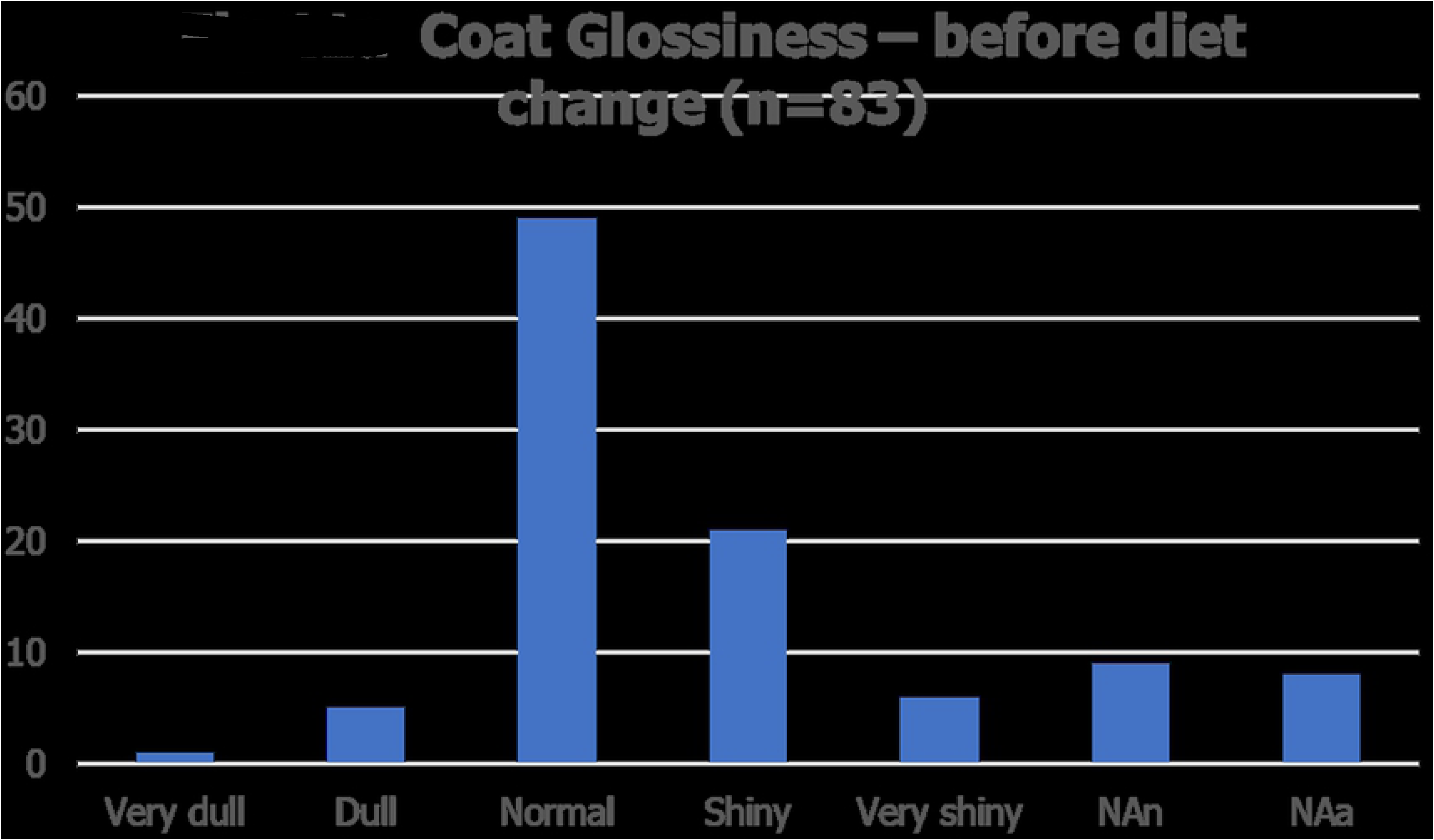
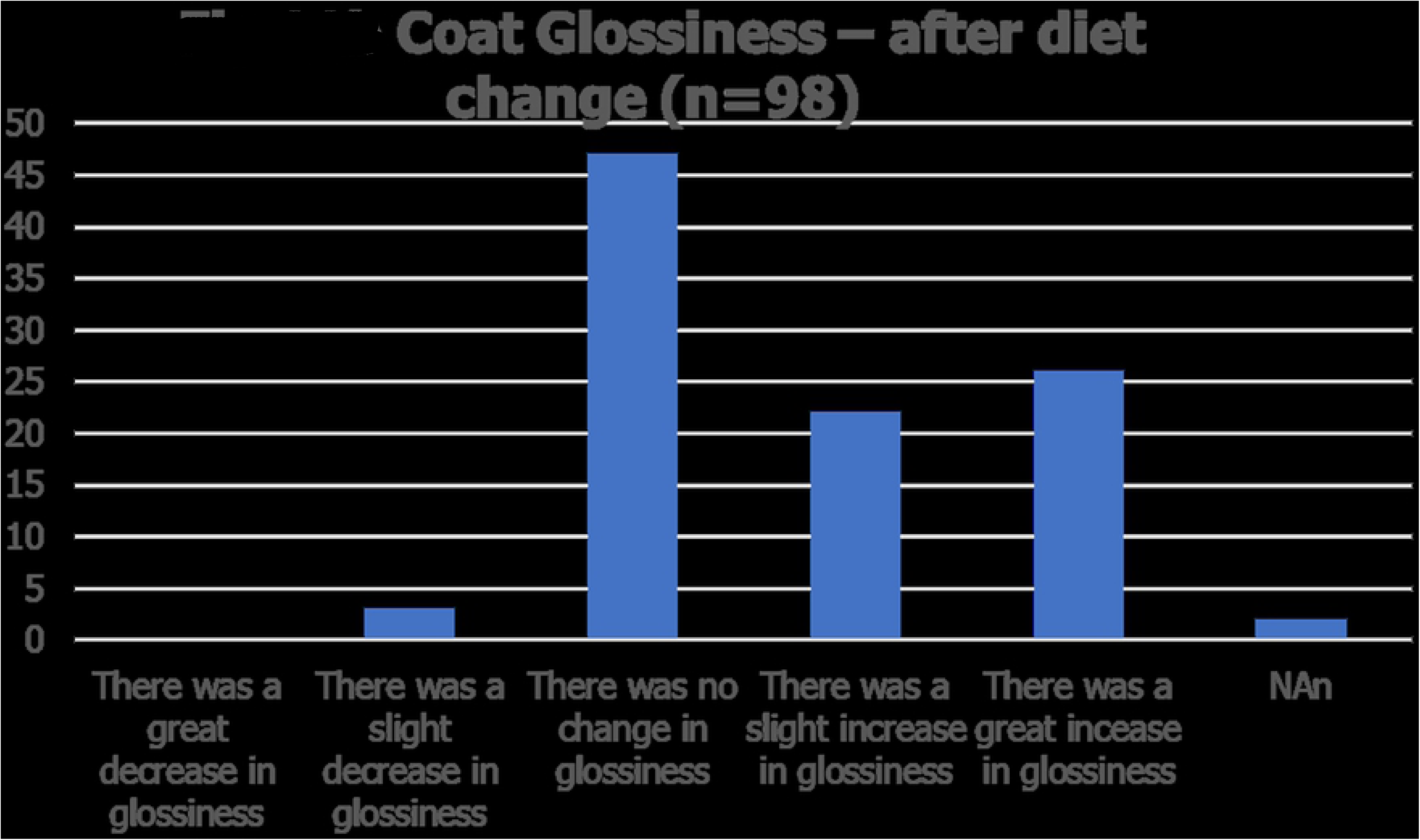

#### Scale (Dandruff)

##### a. Before diet change (n=100)

80 dogs (80.0%) did not have any scale before the switch to plant-based food and 20 dogs (20.0%) did.

##### b. After diet change (n=97) (NAn=3)

One normal dog developed slight scaling after the diet change but there was no change in the other 79 (81.4%). Of the 20 dogs with scale present 3 guardians did not answer the question, of the 17 responses there was no change noted in 4 (23.5%) whereas in the majority of 13 dogs (76.5%%) either a slight improvement (7 dogs – 41.2%) or total resolution of signs (6 dogs – 35.3%) was reported.

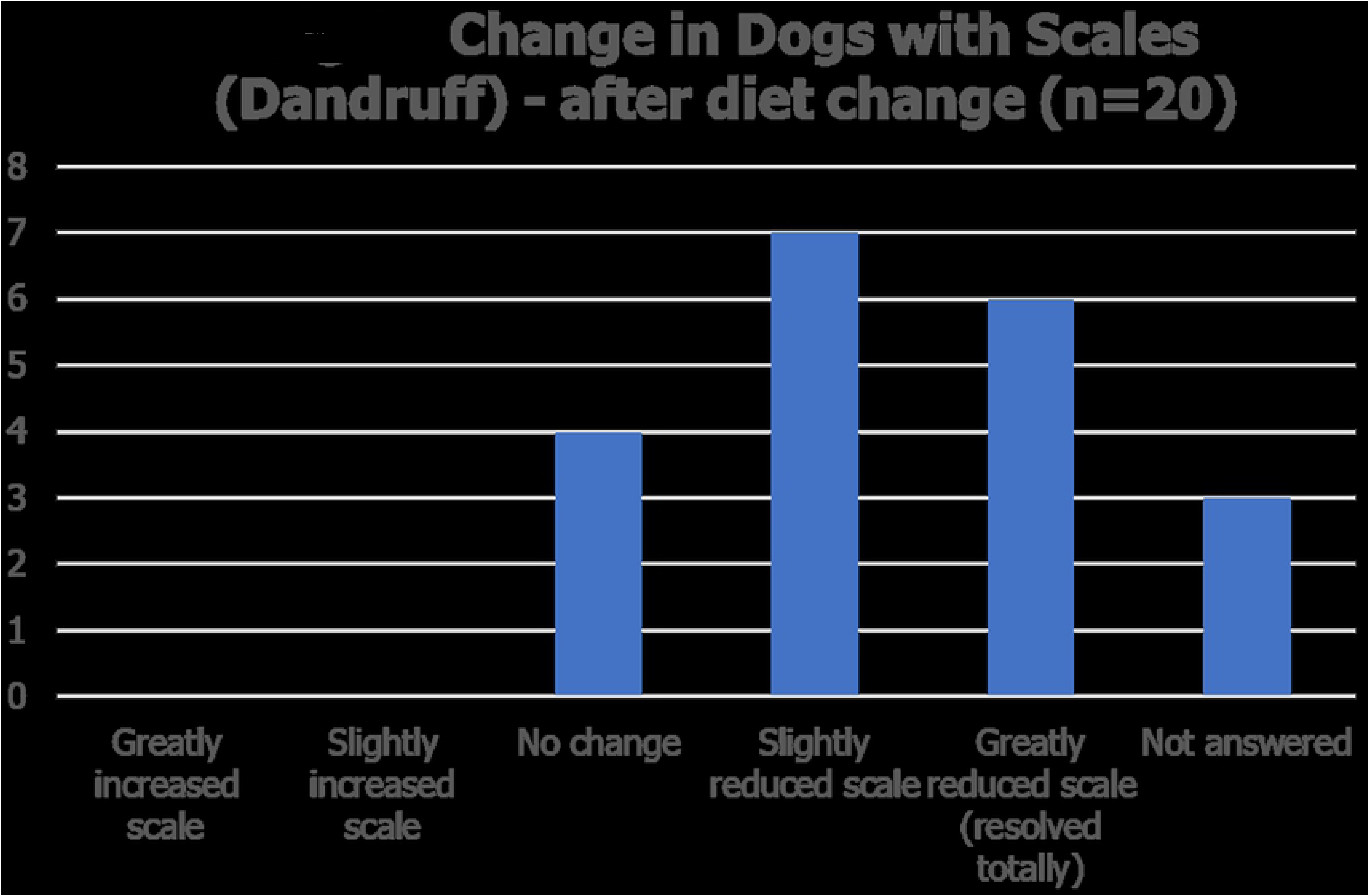

#### Crusting in the ear canal (n=8) (NR=92)

The majority of guardians (92%) reported that their dogs did not have any crusting in their ear canal before switching to the plant-based ration.

Of the 8 dogs that did have crusting in their ears this improved in 7 (85.7%) following diet change, it improved a bit in 5 (62.5%) and resolved totally in 2 (25%). There was no change in the other dog (12.5%). None of the normal dogs developed signs after diet change.

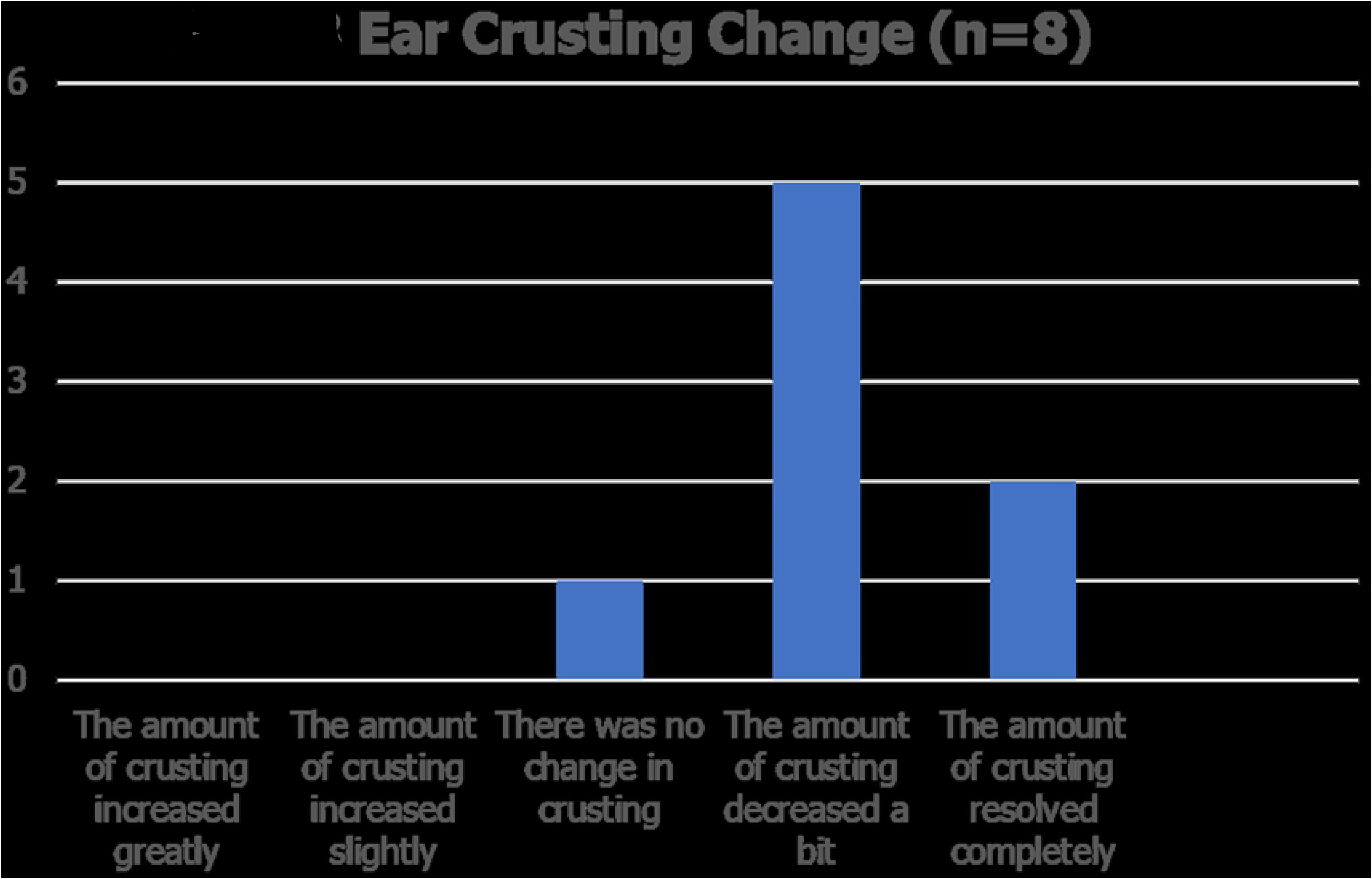

#### Itching (n=71) (NR=29)

Of 100 guardians in the survey 29 (29.0%) reported that their dogs were not itching initially so the majority of 71 dogs (71.0%) were. After changing diet there was no change in 43 (60.6%) but it improved in 37 dogs (52.1%) and in 11 of these (15.5%) the itchiness resolved totally. Itching got slightly worse in 2 dogs (2.8%).

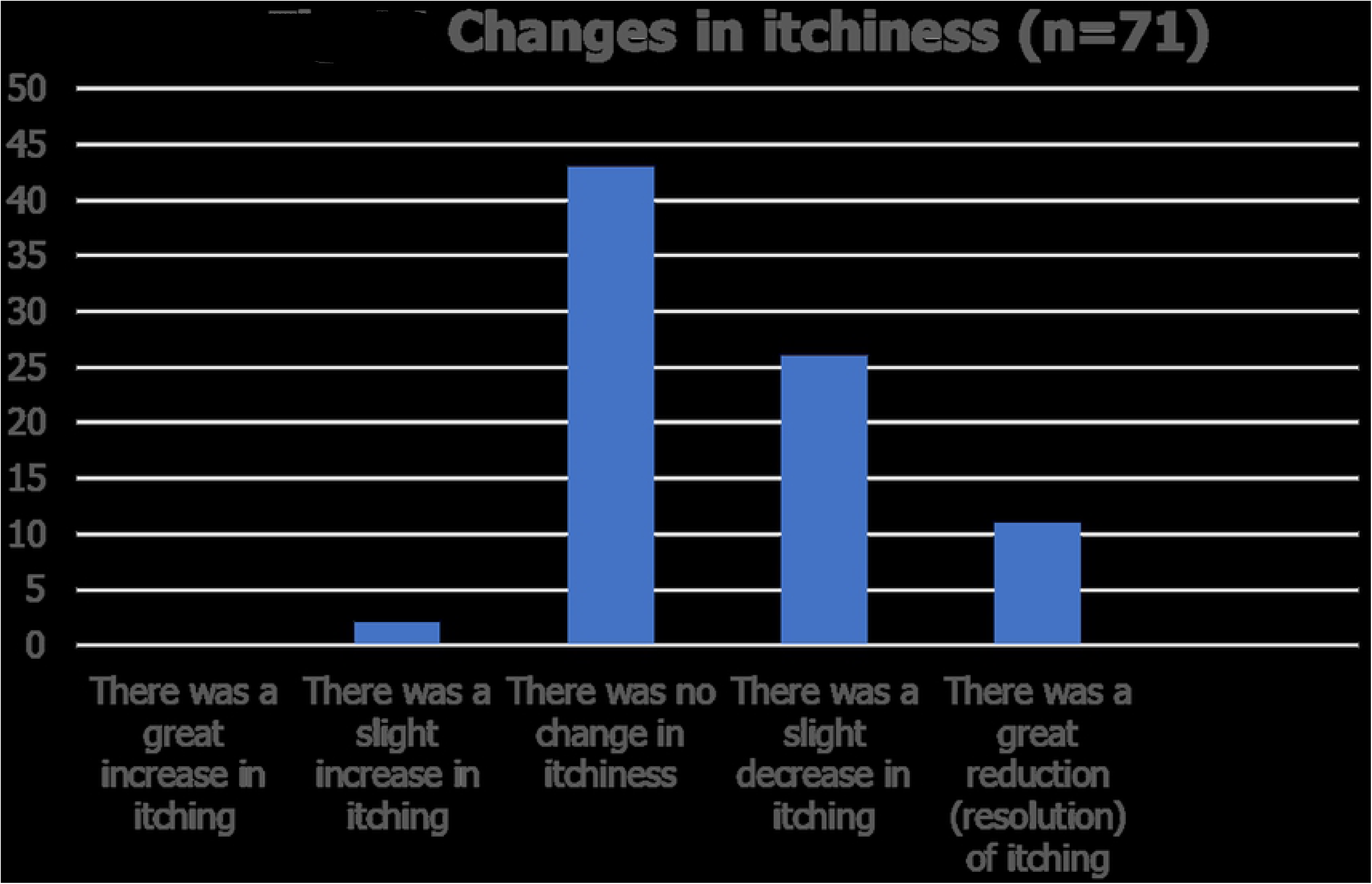

#### Skin redness (erythema) (n=27) (NAn=6) (NR=67)

67 (69.8%) of guardians reported that their dog did not have erythema (reddening) of the skin. Of the 27 (28.7%) dogs that did have reddening there was no change in 15 (55.6%) after switching to the plant-based diet but in 12 dogs (44.4%) the redness improved and in 9 (33.3%) of these dogs there was total resolution of the redness.

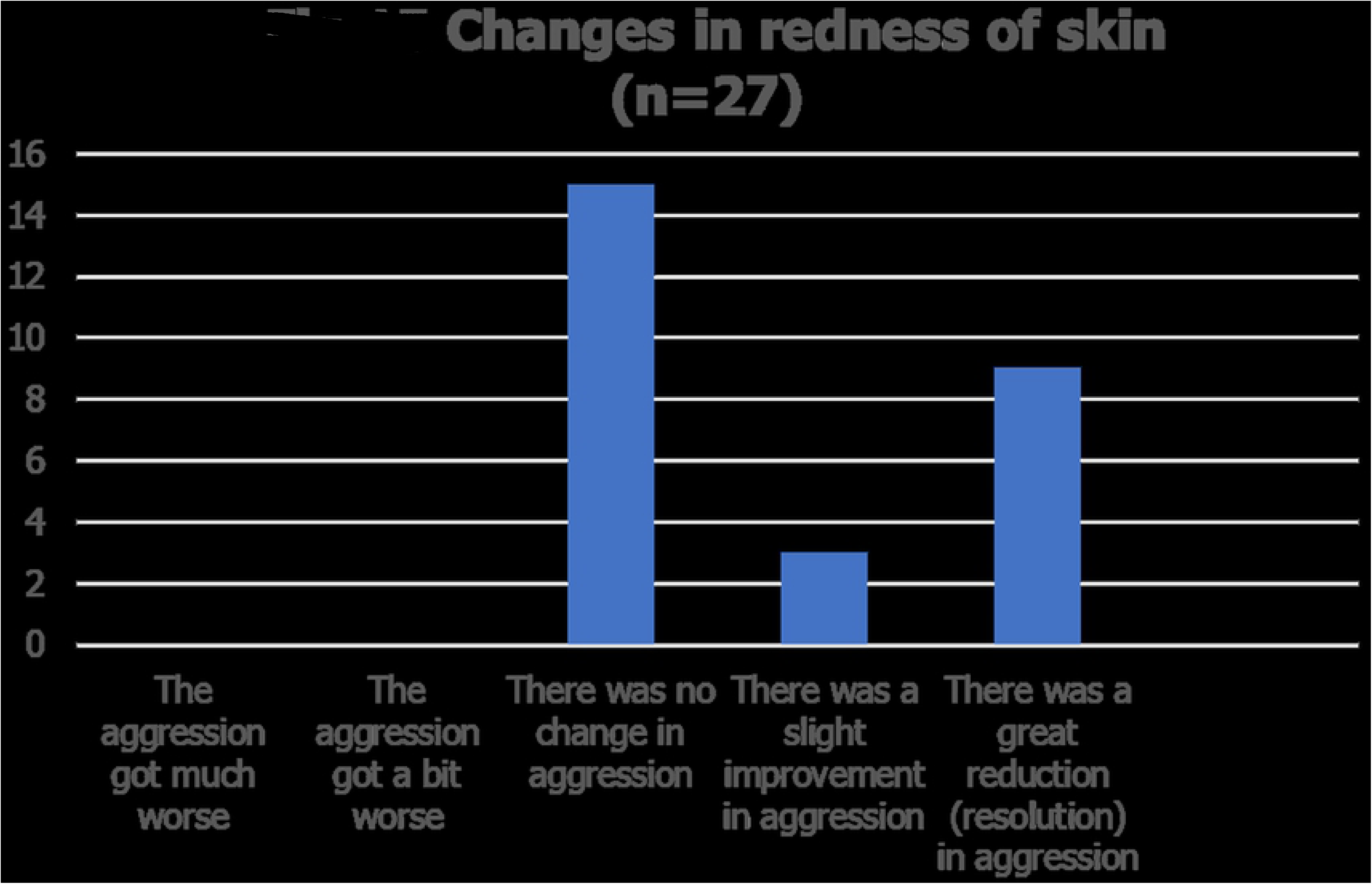

### Behaviour changes

#### Aggression (n=98) (NAn=2)

Of 98 guardians who completed this section 32 (32.7%) described their dogs as showing signs of aggression. After changing to the plant-based food in 23 (71.8%) the aggressive behaviour did not change, in 8 dogs (25.0%) the signs of aggression reduced and in one of these the aggression resolved totally. Aggression got a bit worse in one dog. None of the normal dogs developed aggressive behaviour after switching to the plant-based food.

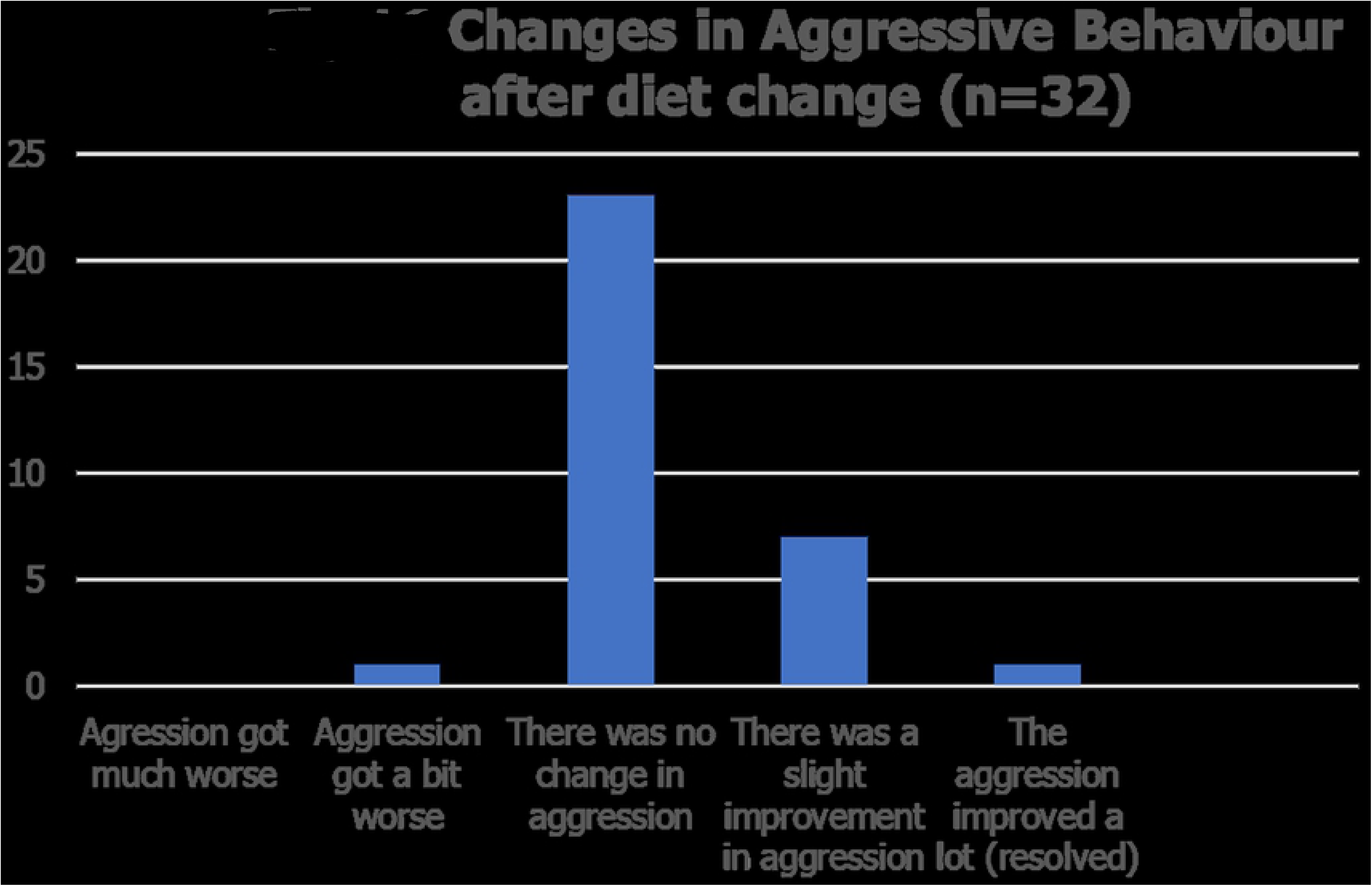

#### Anxiety (n=100)

37 (37%) of dogs were described as showing signs of anxiety prior to changing to the plant-based diet. Afterwards 20 (54.0%) did not show any change in anxious behaviour, 14 dogs improved (*44*.4%) of which 11 (29.7%) improved slightly whilst 3 (8.1%) stopped showing any anxiety. In 3 (8.1%) dogs the anxiety got much worse after changing diet.

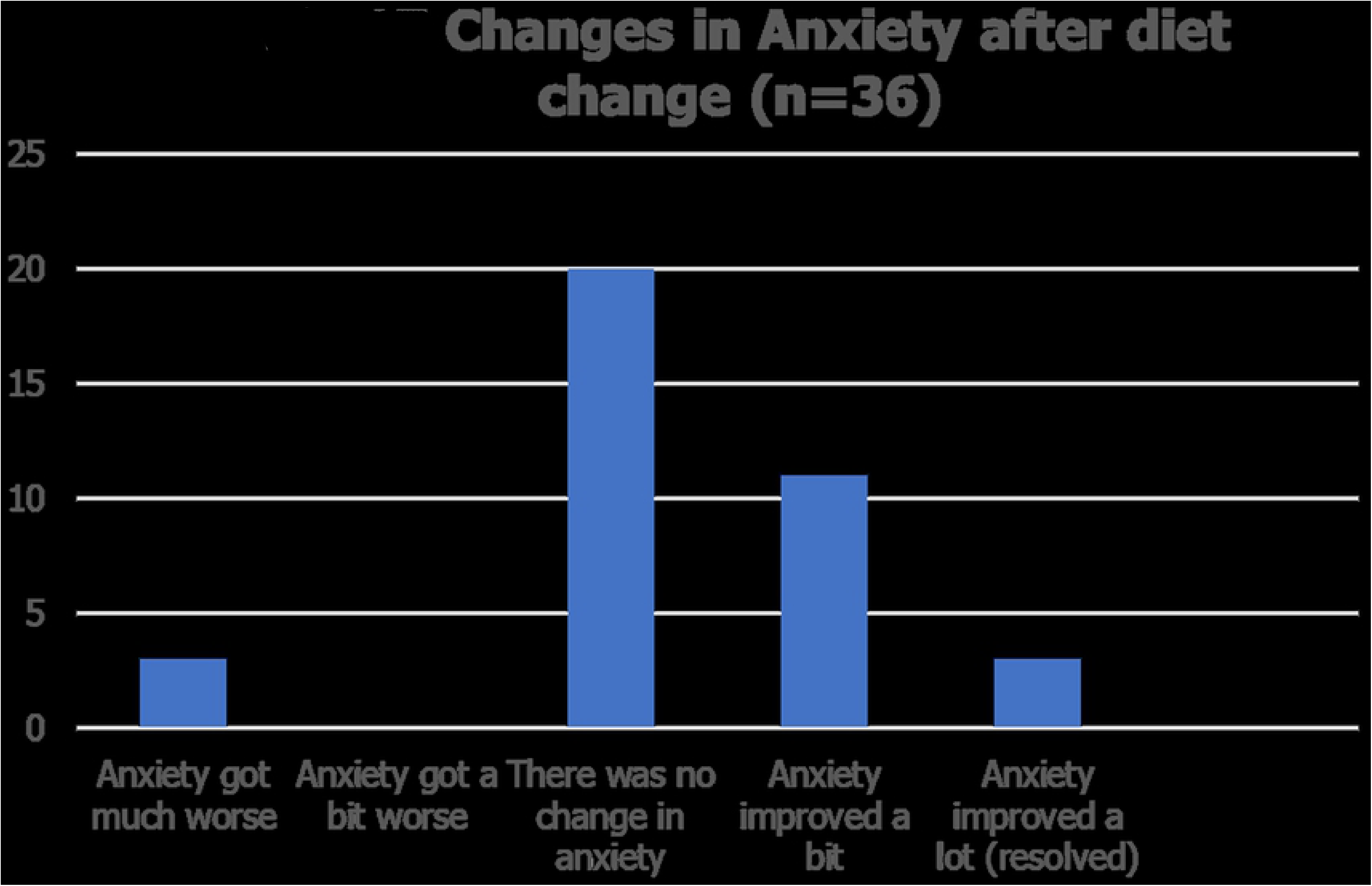

#### Other Behaviours (n=86) (NAn=14)

Of 86 guardians who completed this section only 17 (19.8%) reported that their dog had another behavioural problem before switching to plant-based food and the majority of these (14– 16.3%) were coprophagic. Two dogs had a barking problem, and one each were reported to be showing signs of chewing feet or restlessness

#### Coprophagia (n=14)

After switching in 6 (42.9%) of 14 affected dogs the behaviour resolved totally, one (7.1%) improved slightly, there was no improvement in 6 (42.9%) and one (7.1%) got slightly worse.

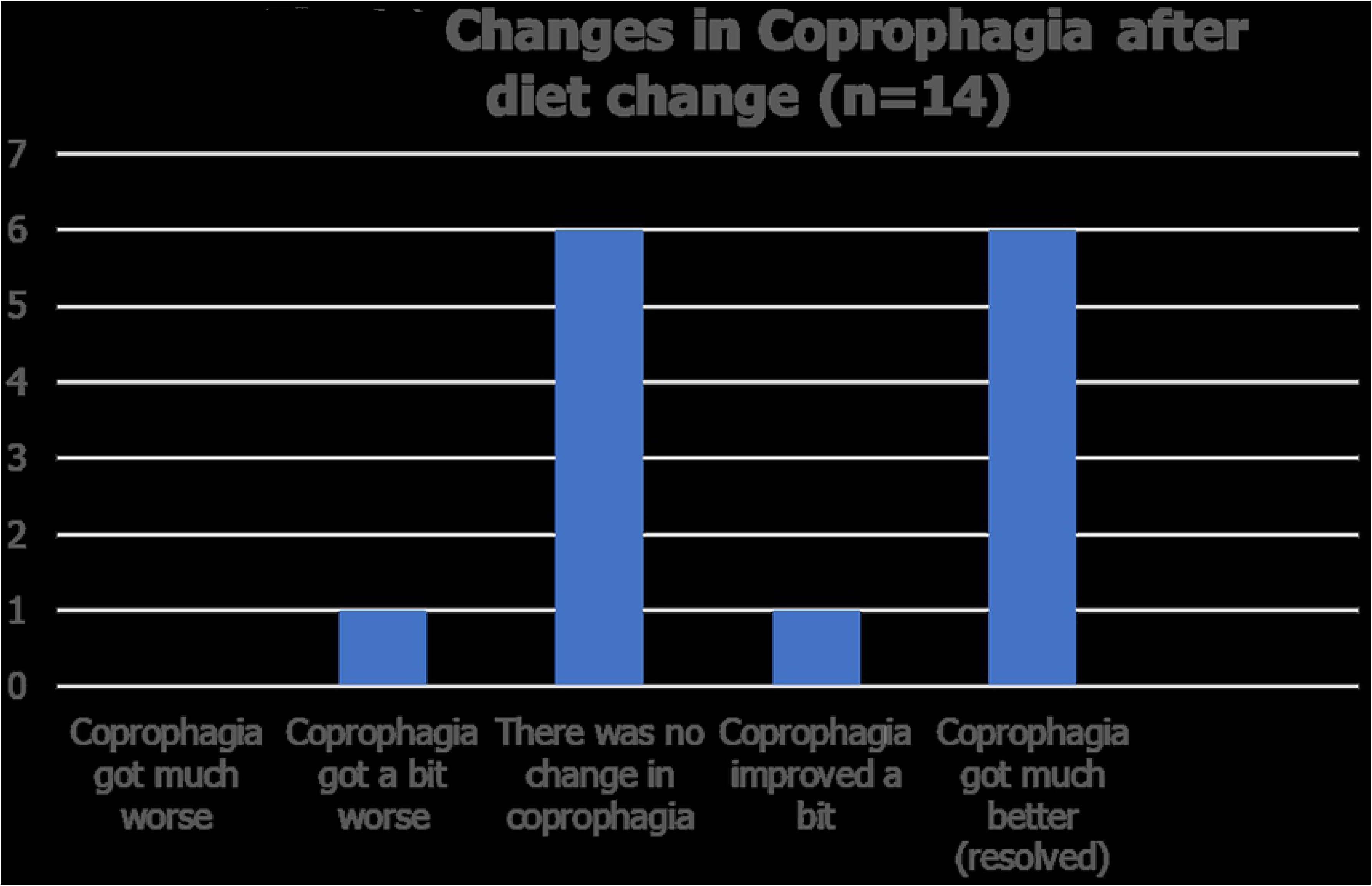

#### Supplementary questionnaire (n=50)

50/100 (50%) of guardians who completed the main questionnaire completed the supplementary questionnaire.

#### Age (n=50)

There was a wide range of ages from 1.5 months to 13 years of age

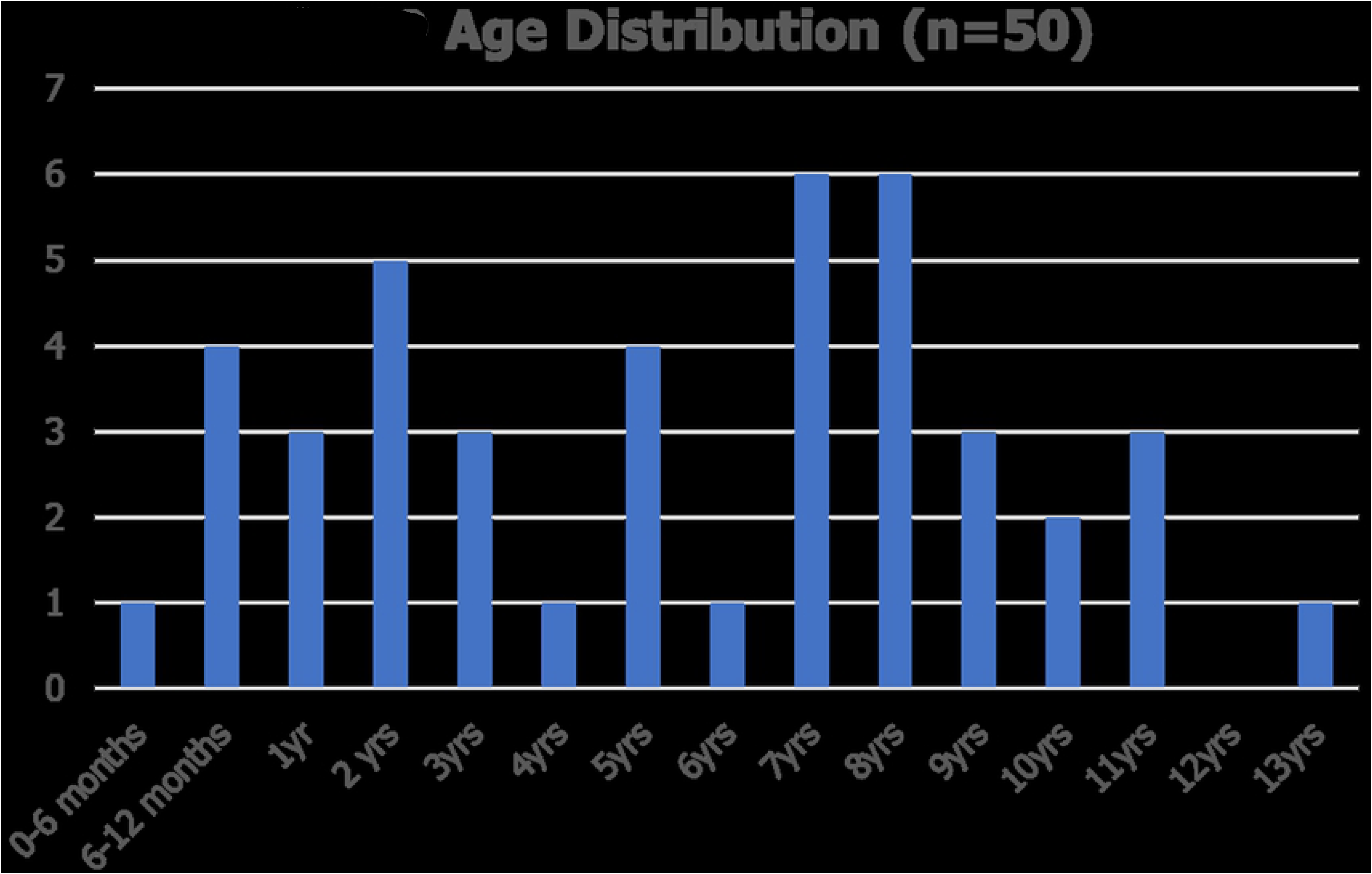

#### Breeds (n=50)

Most dogs (18-36%) were crossbreeds, 4 were Labrador retrievers (8%), there were 3 Cockerpoos (6%) and 2 each of Greyhounds, English Bull Terriers and Hungarian Vizlas. There were single dogs representing 19 other breeds.

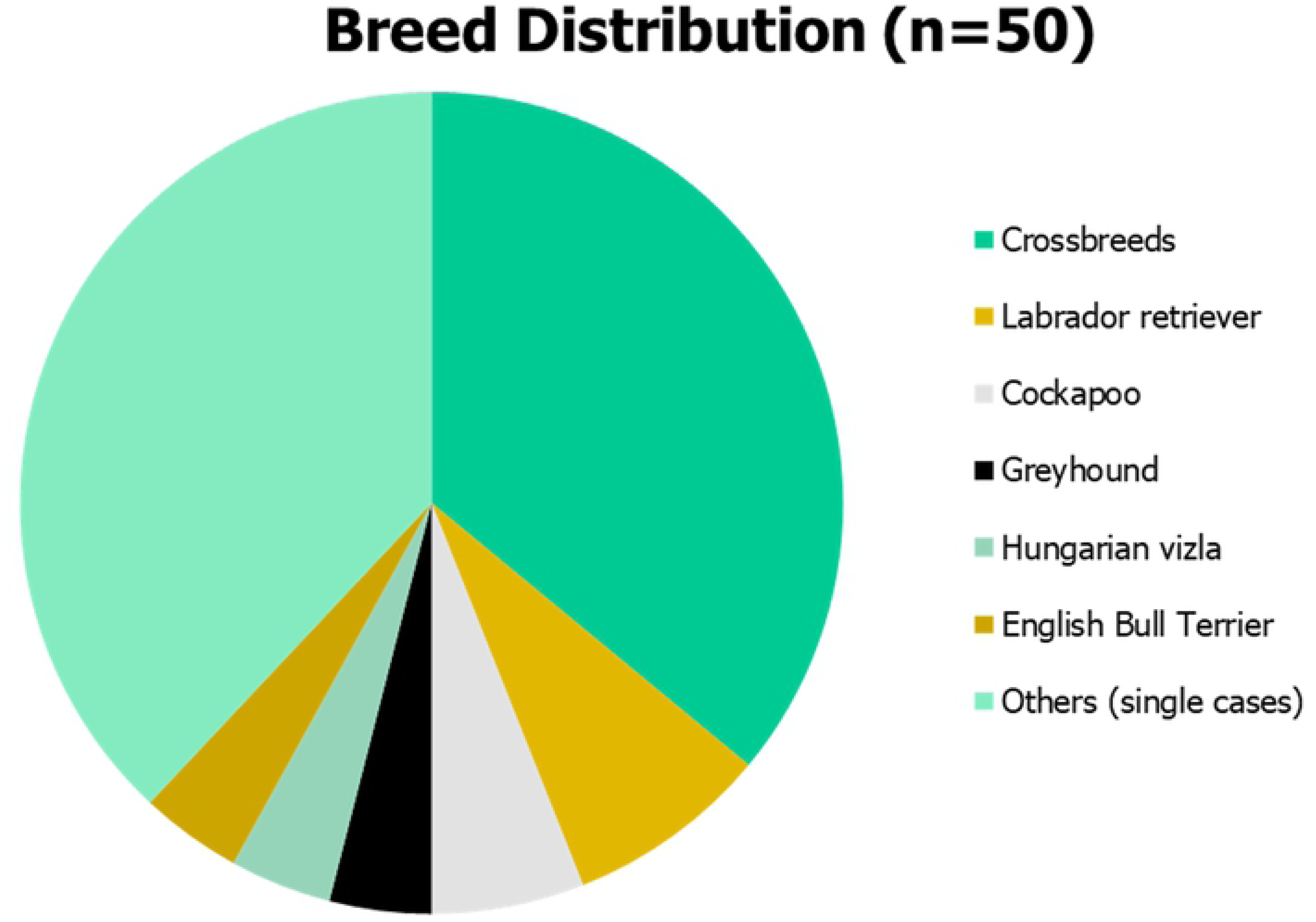

#### Sex (n=50)

There were slightly more males than females Male – 28 (56%); Female 22 (44%)

#### Diet prior to switch to Plant-based food (n=50)

Prior to switching to the plant-based food in this study 11 guardians reported that they were feeding another vegan food (22%) and 39 a meat-based food (78%) of which 4 (8%) were feeding a raw meat ration.

## Discussion

There are a number of important limitations to this study not least that the survey was conducted with a population of dog carers who had decided to feed a vegan food and so were likely to be positive-biased towards recognising positive health benefits.

The survey response rate of 32.6% was in line with reported rates for surveys (Fincham 2008) but was much lower than some scientific Journals now expect which can be as high as 60-80% (Fincham 2008). The non-response bias (67.4%) in this study weakens the reliability and validity of the results but only if one wants to apply the findings of the respondents to the whole population surveyed. The aim of this study was simply to identify whether health benefits were being observed by dog guardians, which it did, and so the non-response bias is less important. A higher response rate could have been achieved by prior communication with potential participants, allowing the survey to be available for a longer period, post-survey follow-ups and reminders, or the deployment of multimodal methods to attract recruits (Fincham 2008).

The dogs in this study had only been fed the food for 3-12 months and so longer follow-up surveys are needed to document long term benefits. Most of the participants were not dog health professionals and so would not be qualified to accurately assess signs of health status. All the outcome measures relating to health status that were provided in the questionnaire, except body weight were subjective not objective, and data collection was incomplete, for example only 50% of respondents to the primary survey provided the additional information requested. So, the results do need to be viewed with a degree of scepticism.

Data derived from pets living under normal home conditions reflect the real-world situation compared to those kept in controlled environments and the overwhelmingly positive results reported here for this specific vegan food, chime with many anecdotal reports of health benefits and provides initial evidence to support further studies.

One concern was that the plant-based diet might not be acceptable (palatable) to dogs, especially if they were used to being fed a meat-based ration which 78% of the dogs were. However, over 82% of the dogs liked the food and ate it all when first given it, and a further 10% ate it all after a short break so 92% of dogs accepted the food immediately. Just 8% of dogs needed to have the food introduced gradually. This is in agreement with another guardian-based study which reported that vegan foods were as palatable as meat-based foods for both dogs and cats (Knight and Satchell 2021).

In dogs many factors affect appetite including odour, taste and texture. Appetite was not affected by the change to plant-based ration in the majority (72.2%) of dogs in this study whereas 16.5% of guardians reported that their dogs’ food intake had increased and some of them (7.2%) were much hungrier and showed begging behaviour for more food. Satiety is induced by the presence of food in the gastrointestinal tract through a combination of neurological (e.g. distension of the tract) and chemical messages (e.g. chemoreceptors in the intestinal wall; absorbed amino acids in the blood) acting on the satiety centre in the brain. Increased food consumption could represent a strong liking for the new food, or a lack of satiety when fed at the recommended feeding rate. Further studies are needed in these dogs to see if the cause was inadequate calorie supply or something else.

The majority of dogs (71%) did not show any fluctuation in body weight which is to be expected if the food was being fed at the appropriate rate to meet the dog’s individual calorie requirements. There are many causes of weight gain or loss in addition to high or low calorie intake and for dogs in which weight varied by more than 10% further investigations would be warranted to determine the underlying reason.

The plant-based food did not affect body weight or body condition score for over 65% of the dogs. Many factors could have caused the increase or decrease in weight and BCS reported for some of the dogs, for example calorie intake or exercise level, and further studies would be needed to determine the reason for the changes in these cases.

Activity levels reportedly increased in 28.9% of dogs after switching to the plant-based food, and in 9.3% a great increase in activity was noted. Only 2 dogs reportedly were less active after changing food, and there are many possible causes for this apart from diet-change that need investigating.

There was no change reported in the frequency of defaecation in the majority (56.0%) of dogs after switching to the plant-based food. There was variability in response in individual dogs with 18 (19.1%) dogs having reduced frequency and 17 dogs (18.1%) having increased frequency. Many factors can influence the frequency of defaecation for example faecal consistency, transit time through the colon, or the presence of dietary fibre and more studies are needed to determine a cause-and-effect relationship with the vegan food which does contain sources of both soluble and insoluble fibres.

The observed effects on faecal consistency were very interesting as 28 out of 32 dogs (87.5%) with soft or watery faeces improved after switching to the plant-based food and in 8 out of 11 dogs with hard or very hard faeces stool consistency improved by becoming softer. Randomised controlled clinical trials would be desirable to establish a clinical role for this vegan food in managing diarrhoea and/or constipation.

Faecal colour can be influenced by many factors including food ingredients and the host microbiome, but also infectious agents and haemorrhage. It was interesting that the majority of guardians who described their dog as having light faeces initially perceived that it had become darker (83%), and at the same time in the majority of dogs with dark brown faeces initially, guardians perception was that it became lighter (88%).

Passing wind is a common problem with some dogs. In this study passing wind reduced in 18 (81.8%) out of 22 dogs reported to be passing a lot before diet change, only 10 guardians (10.4%) reported an increase in flatus after switching even though plant-based ingredients might be expected to increase gas production through the fermentation of fibres

In dogs passing antisocially smelling wind prior to changing diets a switch to the plant-based food resulted in an improvement in 19 out of 26 (73.08%) with a great improvement reported in 17 (65.38%). This vegan food might be beneficial in managing this problem in dogs.

49% of guardians reported that their dogs coat glossiness (shine) improved and 82% of dogs showed an improvement in scaling after switching to the plant-based diet furthermore in over half of the dogs with dandruff the problem resolved totally. There are many factors that can influence glossiness and scaling (dandruff) including nutritional deficiencies: vitamin A, vitamin B complex, zinc and essential fatty acids (Scott and others 1995). So, one of the most likely associations linking improvement after change of diet and clinical signs would be correction of an actual or relative essential nutrient deficiency. Essential n6 fatty acids are important for skin and hair health and plant-based oils provide a higher amount of essential Omega-6 fatty acids than animal-derived fats so dogs switching from meat-based to plant-based rations might experience an improvement

Of the 8 dogs reported to have wax, redness and crusting in their ears, indicative of otitis externa, 2 were being fed another vegan diet before the change and the majority (85.7%) of dogs (including the vegan fed dogs) improved after switching to the study diet. In 2 dogs (25% of those affected) the condition resolved totally. In only one dog there was no change in crusting in the ears. There are many factors that can influence external ear canal condition and these observations are difficult to explain unless the food rectified an underlying dietary deficiency, such as protein (essential amino acids) or fatty acids (Scott and others 1995) that was impairing normal skin physiology or suppressing immune activity. Prospective randomised controlled studies are needed to confirm a cause-and-effect relationship with the diet.

Perhaps surprisingly, 71% of dogs were reportedly scratching before the diet change and the signs improved afterwards in 37 dogs (52.1%) and in 11 of these (15.5%) the itchiness resolved totally. There are many possible causes of pruritus including ectoparasites, bacterial infections, atopy, food sensitivities (allergies or intolerances), hormone imbalance, and dietary deficiencies including Vitamin A, Vitamin B12 (cobalamin), Vitamin D, omega-6 and omega-3 polyunsaturated fatty acid (Saevik and others 2004; Scott and others 1995), and iron (Yonova 2007). Naturally occurring lipid supplementation has been shown to reduce pruritus in dogs (Noli and others 2015).

In humans, pruritus has been associated with low serum zinc and reversed with zinc supplementation and zinc products have been used for many years in people (Sanada 1987; Gupta and others 2014) and dogs (Miller 1989).

It is well documented that compound therapeutic diets can reduce pruritus in dogs (Watson and others 2022; Witzel-Rollins and others 2019), fatty acid supplementation (Bensignor and others 2008) and individual dietary components including Vitamin E (Plevnik Kapun and others 2014) and phytochemicals such as polyphenols may also have anti-pruritic properties in canine atopy (Magrone and Jirillo 2012)

The most likely reasons why a change in diet might result in improvement in itchiness would be correction of an underlying deficiency, removal of a dietary allergen (in this situation animal allergens) or the positive anti-pruritic effect of a nutrient or combination of nutrients in the food. In any future prospective study it would be useful to employ a validated pruritus scoring system for dogs (Rybnícek 2009) against which any improvements can be measured.

Over 28% of dogs had skin erythema (reddening) which indicates inflammation and there are many possible causes of skin erythema including environmental factors, and dietary factors including essential fatty acid, Vitamin E and selenium deficiency (Scott and others 1995). In this study the redness improved in 55.6% of affected dogs and it totally resolved in 12 (44.4%). Further studies are needed to determine the underlying reason for the improvements observed.

Aggression reportedly improved in 25% of dogs following a change in diet and only one showed a slight increase in aggression. In human studies (Werbach 1995) several nutrients have been associated with aggressive behaviour including niacin, pantothenic acid, thiamine, pyridoxine, vitamin C, tryptophan and iron deficiencies. In rodents magnesium deficiency, and manganese toxicity in people also cause aggression.

In dogs, tryptophan and tyrosine, omega-6 and omega-3 fatty acids have been hypothesised to be associated with aggression and other behavioural changes (Bosch 2007). In a thesis and literature review (Harju 2016) canine aggressive behaviour was reported to be decreased by low protein content and tryptophan supplements.

In the study reported here the type of aggressive behaviour being displayed by the dogs was not specified. In a previous study a diet with high protein concentration (32 %) increased fear induced territorial aggression in dogs compared to low (17 %) and medium (25 %) protein concentrations (Dodman et al. 1996). The vegan food in this study is relatively high in protein (30%) and yet aggression reportedly only increased slightly in one dog, but decreased in many more.

In one study plasma concentrations of docosahexaenoic acid (DHA) were lower and the linoleic to α-linolenic acid ratio was higher in aggressive than non-aggressive German Shepherds (Re et al., 2008) suggesting that fatty acids may play a role. In another study dominant aggressive dogs had significantly lower serum concentration of triglycerides than non-aggressive dogs (Pentürk & Yalcin 2003) and both these studies showed that total serum cholesterol and high-density lipoprotein cholesterol levels are lower in aggressive than non-aggressive dogs. Similar associations have been recorded in other species including monkeys, rats and humans (Buydens-Branchey et al., 2000; Hillbrand & Spitz, 1999; Kaplan et al., 1994). This is interesting because the plant-based food in this study contains no cholesterol, although the dog synthesises its own in the liver, and only one dog was reported to have developed a slight increase in aggression.

Anxiety was reported for 36% of dogs however the type of anxiety (e.g. separation or fear) was not specified. In 44.4% of these there was an improvement noticed on the vegan diet and this warrants further studies to determine whether this was a genuine cause-and-effect relationship to the diet Coprophagia is common in dogs and underlying causes include undernutrition, dietary nutrient deficiency, gastrointestinal disorders such as pancreatic exocrine deficiency or inflammatory bowel disease. In this study 46% of dogs stopped this antisocial behaviour after starting the plant-based food which warrants further investigation to see if it might be a useful management strategy.

It should be noted that health benefits were reported in dogs that switched to this vegan diet from other vegan foods, so it cannot be assumed that the positive observations reported for the specific food in this study would necessarily result from feeding other plant-based foods.

## Conclusions

Feedback from 100 dog guardians clearly demonstrates several positive observations of improvements in health in some dogs including in the following areas: faecal consistency, frequency of defaecation, flatus frequency, flatus antisocial smell, coat glossiness (shine), scales on the skin (dandruff), redness of the skin (erythema, inflammation), crusting of the external ear canals (otitis externa), itchiness (scratching; pruritus), anxiety, aggressive behaviour and coprophagia.

These observations could simply be random coincidence relationships and prospective, randomised, controlled clinical studies are needed to confirm the clinical significance of these observations.

Nevertheless, this study confirms several positive health benefits so the hypothesis is disproved, indicating that the feeding of vegan food to dogs does provide health benefits to dogs.

## Acknowledgements

I would like to thank Guy Sandelowsky and Shiv Sivakumar of Omni for allowing me to conduct this survey of their customers with permission to publish the findings regardless of possible negative findings.

## Conflicts of interest

The author is currently a part-time independent consultant to Omni, a part-time external Lecturer at the University of Surrey and Managing Director and Founder of Provet Limited.

Although this study was conducted in collaboration with Omni, members of the company did not influence the process, interpretation or reporting of the data analysis.

## Notes

### Competing Interest Statement

The author was working as an independent consultant specialist in small animal clinical nutrition to Omni at the time of this study.

